# Quantifying Adaptive Evolution of the Human Immune Cell Landscape

**DOI:** 10.1101/2023.10.06.559946

**Authors:** Irepan Salvador-Martínez, Jesus Murga-Moreno, Juan C. Nieto, Clara Alsinet, David Enard, Holger Heyn

**Affiliations:** CNAG, Centro Nacional de Análisis Genόmico, Barcelona, Spain; Departament de Genètica i de Microbiologia, Universitat Autònoma de Barcelona, Barcelona, Spain; University of Arizona Department of Ecology and Evolutionary Biology, Tucson, United States

**Keywords:** **Keywords.** Adaptation, Immune System, Human Cell Atlas

## Abstract

The human immune system is under constant evolutionary pressure, primarily through its role as first line of defence against pathogens. Accordingly, population genomics studies have shown that immune-related genes have a high rate of adaptive evolution. These studies, however, are mainly based on protein-coding genes without cellular context, leaving the adaptive role of cell types and states uncharted. Inferring the rate of protein-coding genes adaptation in developing and adult immune cells at cellular resolution, we found cell types from both the lymphoid and myeloid compartments to harbour significantly increased adaptation rates. Specific cell states, such as foetal Pre-Pro B cells and adult T resident memory CD8+ cells show highly elevated rates of adaptation. We further analysed activated cell states, specifically, iPSC-derived macrophages responding to various challenges, including pro- and anti-inflammatory cytokines or bacterial and viral infections, the latter simulating the evolutionary arms race between humans and pathogens. Here, we found positive selection to be concentrated in early immune responses, suggesting benefits for the host to adapt to early stages of infection to control pathogen numbers and spread. Together, our study reveals spatio-temporal and functional biases in human immune populations with evidence of rapid adaptive evolution and provides a retrospect of forces that shaped the complexity, architecture, and function of the human body.

## Introduction

The need to respond to ever-changing challenges posed by pathogens makes host-pathogen interactions a clear example of evolutionary arms race. In the last couple of decades, many population genetics and molecular evolution studies, which study the change in genetic composition of populations and species, have shown that the immune system is under constant evolutionary pressure and provided multiple examples of genes evolving under increased rates of adaptation. The evolution of human immune genes has even been associated with specific diseases and pathogens, such as malaria[1,2] and the Black Death pandemic that killed 30-50% of the Afro-Eurasian population in the XIV century[3]. At a deeper evolutionary time scale, studies on primates and mammals showed that a significant amount of adaptation is due to the arms race between viruses and their hosts[4]. Such a competition not only plays a selective pressure leading to the rapid fixation of advantageous amino acid changes on multiple antiviral proteins-coding genes, but also in non-antiviral protein-coding genes. As an example, members of the APOBEC family, PKR and TRIM5, antiviral molecules that directly attack viral molecules, as well as TFRC and ANPEP, non-antiviral proteins involved in iron transportation and viral receptors respectively, are well-known examples with unusual rates of adaptation in several organisms including humans[4–9]. However, previous knowledge of specialised immune genes is required to make the connection between the host’s positive selection process and pathogens, and such well understood examples represent only a small fraction of all adaptation events in immune genes[4].

Technically, previous studies on immune gene evolution have focused on measuring *ω* (the ratio of non-synonymous, *d_N_*, to synonymous, *d_S_*, substitution per site)[10–12], to determine pervasive or episodic adaptive evolution of the immune system[4–7,9]. Nonetheless, non-synonymous changes may alter the function and protein-coding structure, so they tend to be deleterious and removed from the population by the action of purifying selection, which restricts the ability of *ω* to identify positive selection and tends to mask the impact of adaptive substitutions. To circumvent these limitations, the McDonald Kreitman test (MK-test[13]) combines both divergence and polymorphic data to summarise the rate of adaptation with the quantity *α* (the proportion of adaptive non-synonymous substitution)[14]. Because adaptive mutations are supposed to contribute almost exclusively to divergence and not polymorphism, polymorphic data is used as a background to calibrate the expected divergence rate under the neutral model in the presence of purifying selection[15,16], increasing the power of detecting positive selection. However, during the last decades, it has been repeatedly shown that weakly selected polymorphism can attain low and high frequencies[17–20], depending on the underlying Distribution of Fitness Effects (DFE)[21], biasing MK-test and *α* inferences. To account for these biases, we used ABC-MK, an MK-test extension proposed by Uricchio et al.[22], Murga-Moreno et al.[23]. This approach is particularly useful when weakly advantageous alleles segregate along with the evolutionary process known as background selection (BGS). BGS decreases neutral genetic diversity when neutral variants are removed together with the deleterious variants they are linked to. BGS can also impact positive selection signatures, especially at high frequencies[22]. ABC-MK is an Approximate Bayesian Computation (ABC) which takes advantage of *α*_(_*_x_*_)_ summary statistics proposed by Messer and Petrov[24]. By modelling *α*_(_*_x_*_)_ with multiple DFE random parameter combinations along with BGS, ABC-MK exploits *α*_(_*_x_*_)_ patterns to infer *α* while providing flexibility to analyse heterogeneous datasets from the human genome, where BGS strength is variable and weakly adaptive variants are an essential component of adaptation[22,23]. In addition to quantifying adaptation with *α*, we also estimate *ω_a_*, the rate of adaptive non-synonymous substitution relative to the mutation rate, and we explore the relationship between adaptation and evolutionary gene age in the human immune system by a gene phylostratigraphy approach. This method involved ranking genes into different Phylostrata based on their evolutionary emergence time[25].

Other important challenges, when studying adaptation in the immune system, are the spatio-temporal dynamics and cell type specificity of immune gene expression that have not been taken into account in previous studies, mainly due to a lack of comprehensive transcriptomic reference maps of the immune system at the cell type level. In recent years, the Human Cell Atlas (HCA) has created comprehensive transcriptome reference maps of healthy human organs and tissues at different developmental and adult stages, with the ultimate objective of characterising every cell type in the human body[26]. As part of this effort, the Human Developmental Cell Atlas bionetwork generated a reference map of the developing immune system[27] and a complementary study provided an adult multi-tissue immune cell atlas[28]. Together, both atlases represent the most comprehensive transcriptomic characterization of the human immune system and provide the basis for a better understanding of immune protein adaptation at cell type resolution.

In here, we used ABC-MK along with the developmental and adult immune cell atlases, to quantify adaptation at the cell type level since humans split from chimpanzees. This helped us to understand immune protein adaptation within human immune cells and to identify immunological challenges driving immune adaptation. By leveraging spatio-temporal single-cell resolved reference maps of *>* 1, 2 million cells, we pinpoint immune cell types and states under high adaptation rate. To better understand the selective pressures, immune response timing and the role of natural selection on human immune cells, we analysed the innate immune response to different stimuli that mimic various inflammatory and pathogen exposures, using activated induced pluripotent stem cell (iPSC)-derived macrophages, a successful pathogen infection model. Altogether, our analyses of adaptive evolution charted the landscape of cell types contributing to the arms race between the human host and its pathogens and helped to clarify the origin of abundant host immune protein adaptation, allowing us to hypothesise immune functions and pathogen classes that drove immune adaptation during human evolution.

## Results

### Building an Immune Atlas of Adaptation

The Human Developmental Cell Atlas bionetwork generated a reference map of the developing immune system with *∼*900, 000 cells from 25 donors and more than 100 annotated cell states. The developmental atlas includes data from haematopoietic organs (yolk sac, liver, bone marrow), lymphoid organs (thymus, spleen and lymph node), non-lymphoid tissues (skin, kidney and gut) and spans 4 to 17 weeks after conception[27]. The adult multi-tissue immune cell atlas comprised *∼*300, 000 cells from 21 donors and 16 tissues. Selected tissues included primary (bone marrow) and secondary (spleen and lung-draining and mesenteric lymph nodes) lymphoid organs, mucosal tissues (gut and lung), as well as blood and liver[28]. In the Developmental Immune Cell atlas the immune compartment represented hematopoietic stem cell (HSC) and hematopoietic progenitors, erythro-megakaryocytes as well as the major myeloid and lymphoid cell lineage populations. The spatio-temporally resolved reference captured the emergence of immune lineages from hematopoietic progenitors, characterising immune cell development before tissue seeding to the developing lymphoid organs, such as the thymus, spleen, lymph nodes, and peripheral non-lymphoid organs. On the other side, the Adult Immune Tissue atlas provided the characterisation of mature immune cells in circulation and specialised to tissue microenvironments, such as tissue-resident macrophages and memory T cells. Together, both datasets provide an annotated resource that represents a complete diversity map of human immune cells in steady state from the embryo to the adult.

In order to identify human immune cell types with signatures of selection in development and adulthood and to generate a comprehensive Immune Atlas of Adaptation (Figure 1), we retrieved the annotated single-cell datasets from both resources and de novo computed cell type-specific differentially expressed (DE) genes. Specifically, after removing cell clusters reported as cycling (no cell type specificity), cell doublets or low quality cells, we performed a differential gene expression test for each cell population compared to all remaining cells in the same compartment (Wilcoxon test, p-value *<* 1 *·* 10*^−^*^4^, logFC *>* 1). For each cell population, we used the top 500 DE genes sorted by logFC value (or all DE genes for populations with *<* 500 genes) to normalise differences in the number of differentially expressed genes between cell populations, while maximising the number of genes to perform accurate ABC-MK estimations in human lineage following Murga-Moreno et al.[29] results. In the developmental atlas, some cell types from the progenitors compartment are also present in an additional compartment. For example, Pre-ProB cells and Promonocytes are also part of the Lymphoid and Myeloid compartments, respectively. For these cell types, as the DE genes were obtained independently for each compartment, we report the ABC-MK test results obtained for each compartment.

**Figure 1.**
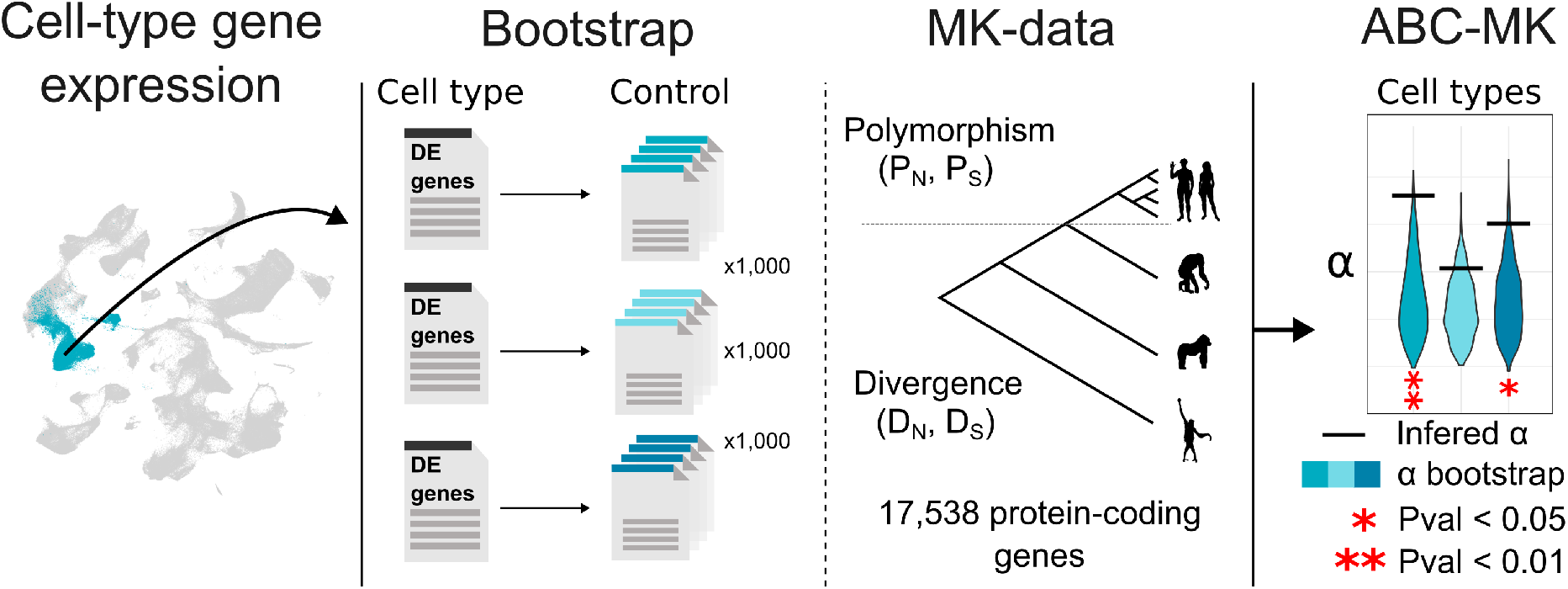
Methodological abstract. Single-cell transcriptomic datasets from the Human Developmental Immune Cell Atlas[27] and the adult multi-tissue immune cell atlas[28] were retrieved and the top 500 differential expressed (DE) genes for each immune cell type was computed de novo. For significance tests, we compare each dataset with 1000 control datasets having similar average values of multiple confounding factors (see Methods). We ran ABC-MK to measure the adaptation rate using data from the Thousand Genomes Project[35] as polymorphic data and human reciprocal orthologs on chimpanzee, gorilla and orangutan genome assemblies as divergent data.

To determine the proportion of adaptive non-synonymous substitution (*α*) driven by weakly and strongly beneficial alleles (*α* = *α_S_*+ *α_W_*), we ran ABC-MK independently for each cell population[22]. Each estimation used random parameter combinations to estimate 10^5^ different *α*_(_*_x_*_)_ combinations accounting for multiple Distributions of Fitness Effects (DFE) and background selection, while sampling empirical data and accounting for partial recombination (see Methods, Supplementary Table **??**). We used ABCreg[30] to perform ABC comparing ABC-MK sampled *α*_(_*_x_*_)_ and empirical datasets. Because *α* prior values range from 0 to 1 to measure the proportion of adaptive substitutions, we used the mode of the posterior distribution to quantify *α*, *α_S_* and *α_W_*, better accounting for cases where adaptation is low or absent (posteriors distributions displaced to 0 values). To measure a significant increase in adaptation, we build a null distribution accounting for 1,000 control datasets matching the size of the analysed immune genes subset for each cell population, following a bootstrapping approach[31,32]. The procedure allowed us to match, resample and compare genes with similar overall average values for many confounding factors that could affect the adaptation rate and mislead the comparison (see Methods for further information)[31,33,34]. We evaluated each cell population estimation in accordance with its corresponding null empirical distribution. Thereby, we ensured statistically significant outcomes surpassing the empirical distribution to arise primarily from positive selection related to the cell populations’ immune function, rather than other confounding factors (see Methods for a detailed list of possible confounders), providing an unbiased estimation of the false discovery rate for adaptation within each dataset. In addition, we replicated the analyses by estimating *ω_a_*, the rate of adaptive non-synonymous substitution relative to the mutation rate, to provide further evidence of protein adaptation (see Supplementary Figures **??**,**??**,**??** and Supplementary Tables **??**,**??**,**??**). Our results showed a picture of highly elevated adaptation, similarly on *α* and *ω_a_* estimations, in diverse immune cell populations in both the developing embryo and in adult human tissues.

### Most foetal progenitor cell types have a high adaptation rate

In the Developmental Immune Cell atlas, we found eight hematopoietic stem and progenitor states, nine lymphoid, three myeloid and one Megakaryocyte population with significantly elevated adaptation rates (Figure 2, Supplementary Figure **??**, Supplementary Table **??**, **??**). Intriguingly, we found Hematopoietic Stem Cells (HSCs) and seven Hematopoietic Progenitor Cells (HPCs) with a high adaptation rate in the progenitor compartment. Specifically, five out of seven myeloid progenitors (MEMP, CMP, GMP, Promonocytes and early Megakaryocytes) and two out of four lymphoid progenitors (LMPP and PreProB) showed high adaptation rate. HSCs give rise to all cell lineages of the immune system and are the only hematopoietic cells with self-renewal and full differentiation capacity through lineage progenitor stages (e.g. Lymphomyeloid primed progenitor and Megakaryocyte–erythroid progenitor, LMPP and MEMP). HSCs are, however, also a sensory hub of the immune system, sensing and responding to molecular cues and physical interactions with their niche’s complex multicellular network[36]. For example, depletion of certain blood cell populations activates HSCs and subsequently HPCs to undergo differentiation[37]. The adequate readout of signals from the hematopoietic stem and progenitor niche is therefore crucial to ensure haematopoietic homeostasis in development and throughout life[36]. Additionally, HSCs respond to infections and inflammatory signals through different mechanisms, including sensing pathogen-derived products and proinflammatory cytokines produced after pathogen infection[37]. Considering the above, we hypothesise that the high adaptation rate inferred in foetal HSCs and HPCs is related to their key position as a guardian of the hierarchy controlling homeostasis and immunomodulation in the developing embryo.

**Figure 2.**
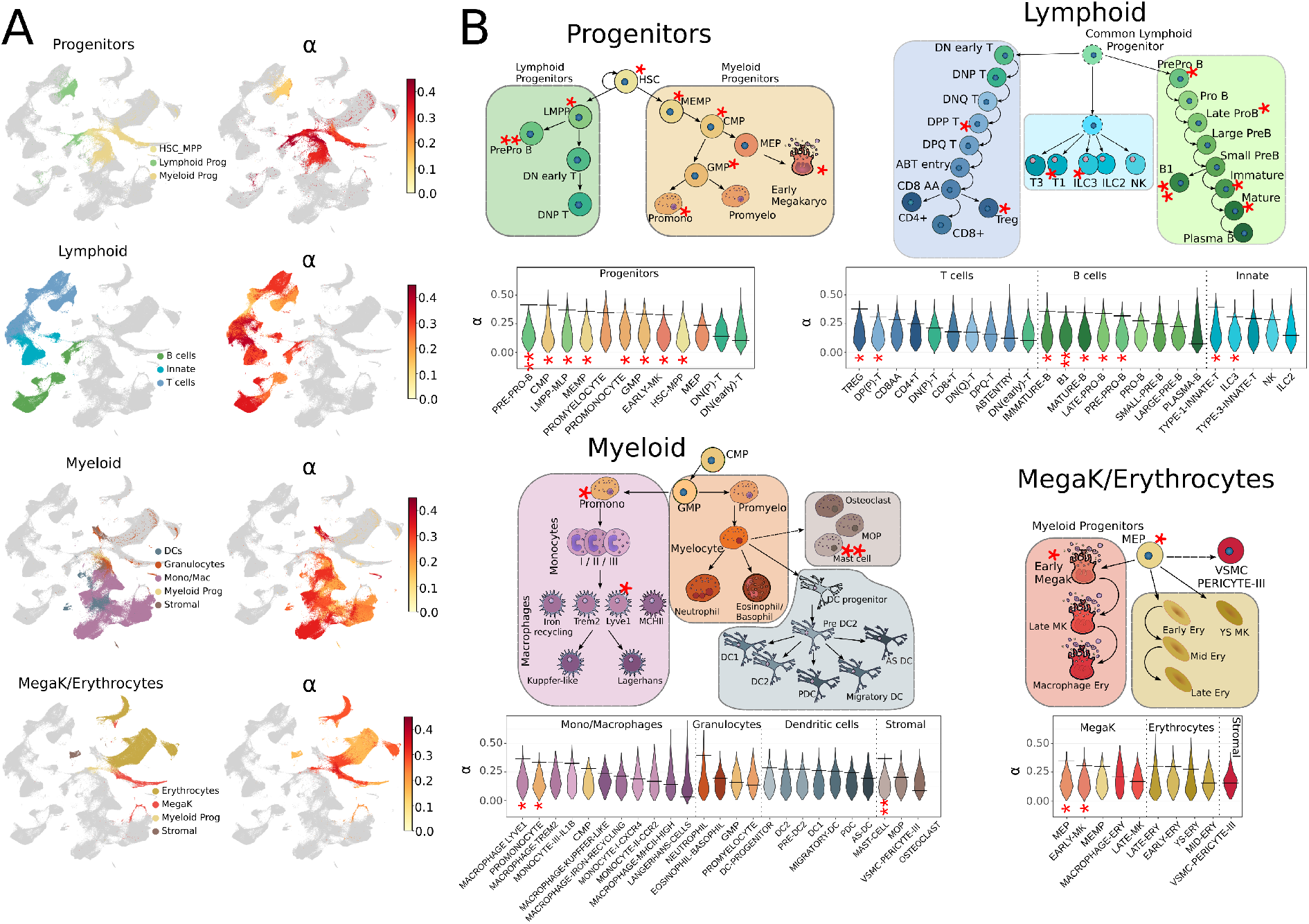
Protein adaptation in the Developmental Atlas. A) On the left, each compartment is highlighted in the global UMAP (from[27]) with a colour code that corresponds to specific lineages. On the right, the inferred *α* value for each compartment is projected on the same UMAP. Note that the *α* value is inferred at the cell-type level. B) A scheme with the putative lineage relationships between the cell types within each compartment is shown. Lymphoid cells with a dashed outline are not part of the analysed dataset and are only depicted to clarify lineage relationships. Under each scheme we show the inferred *α* value of each cell type as a horizontal bar and the *α* value distribution of its control bootstrap dataset (*n* = 1000) as a violin plot. The asterisks highlight the cell types with a significant high *α* (* p-value *<* 0.05, ** p-value *<* 0.01).

### Tregs, ILC3, B1 and B cells under maturation among foetal lymphoid cells with high adaptation rate

The lymphoid compartment is subdivided into three lineages: T cells, B cells and Innate lymphoid cells. T cells and B cells are the major components of the adaptive immune system in humans. Noteworthy, adaptive immunity is a shared feature of all vertebrates with jaws (gnathostomes) and thought to have evolved more than 400 million years ago in the placoderms, the first jawed vertebrates[38]. T and B cells use cell surface receptors to recognise antigens of pathogens that trigger cytotoxic activation and the production of antibodies[39]. Two additional lymphoid cell types are devoid of antigen receptors: Natural Killer (NK) cells and Innate lymphoid cells (ILCs). Functionally, NK cells complement cytotoxic T cells in killing infected cells, whereas ILCs share many functions of T cells, such as sensing stress signals, microbial compounds and the tissue cytokine milieu[40]. Consistent with the finding in the haematopoietic stem cell and progenitor compartment, significant signals of adaptation were found in lymphoid progenitor states. Specifically, we found five out of nine lymphoid cell subtypes in the B lineage (Pre Pro B, Late ProB, B1, Immature and Mature B cells), two out of ten in the T cell lineage (DPP and Treg) and two out of five in the Innate lymphoid cell subtypes (ILC3 and Type 1 Innate T) with a high adaptation rate. From the spatio-temporal perspective, the high adaptation of Immature and Mature B cells coincides with the migration of Immature B cells in the foetal bone marrow or liver to the spleen, where immature cells transition to mature B cells. This transition is evident in the observed organ distribution of annotated cell types in the Developmental Atlas, as the proportion of Immature and Mature B cells in the liver decreases from 36% to 6%, whereas it increases in the spleen from 43% to 81% (Supplementary Figure **??**). During this maturation process, a checkpoint is reached where potentially autoreactive B cells are eliminated[41]. Failure to eliminate autoreactive cells can lead to the development of autoimmune diseases later in life, such as Systemic Lupus Erythematosus and Rheumatoid Arthritis[41]. Similarly, regulatory T cells (Treg) control the immune response and help prevent autoimmune diseases[42]. In contrast, the function of B1 cells and innate lymphoid cells is to provide protection from external challenges. B1 cells are a subclass of B cells that are involved in the humoral immune response and have been exclusively described during embryo development[43,44]. Although B1 cells are not part of the adaptive immune system (lack memory response), they perform similar roles as other B cells, such as the production of antibodies against antigens and acting as antigen-presenting cells. On the other hand, ILC3 participates in innate mechanisms on mucous membranes, contributing to host-commensal mutualism and pathogen clearance. They are also involved in the activation of NK cells, including cell lysis and secretion of pro-inflammatory cytokines, and protect the intestinal mucosa from infections of various pathogens[40].

### Lyve1+ macrophages and Mast cells among foetal myeloid cells with high adaptation rate

Innate immunity, as displayed by myeloid cells, is a more ancient form of host defence against infection that involves pattern recognition systems (e.g. RLRs and TLRs), whose molecular components are shared by all multicellular animals[45]. The myeloid compartment is subdivided in four main lineages: Monocytes/Macrophages, Granulocytes, Dendritic cells and Stromal cells (Figure 2). Myeloid cells are highly plastic, i.e. changing cellular states depending on external stimuli, and are recruited to tissues after pathogen invasion via chemokine receptors. Their role, however, is not limited to immune responses; myeloid cells are known to play an important role in supporting tissue homeostasis and development[46]. In the myeloid compartment we found two out of eleven cell states in the Monocyte/Macrophage lineage (Macrophage Lyve1 High and MEMP) and one out of three cell types in the stromal compartment (Mast cell) with significantly elevated adaptation rate (Figure 2B).

Macrophages with high Lyve1 expression have been found to localise in a specific niche surrounding blood vessels[47] and to have an important role maintaining homeostatic vascular tone by regulating collagen production[48]. Suo et al.[27] showed that the Lyve1 subpopulation of macrophages in the yolk sac exhibits an increased self-renewal potential across various organs. Mast cells, with a long history predating the development of adaptive immunity, are primarily renowned as significant sources of mediators responsible for acute allergic reactions and IgE-dependent immediate hypersensitivity reactions[49]. Furthermore, both early macrophages and Mast cells were noted for their roles in processes such as angiogenesis, tissue morphogenesis, and the maintenance of tissue homeostasis before adopting their more traditional immunological functions. We therefore hypothesise that the functions under adaptation in Mast cells and Lyve1 macrophages are related to their role in tissue support and homeostasis.

Finally, in the Megakaryocyte and Erythroid compartment, we only found two out of ten cell types with high adaptation rate: early megakaryocytes and MEP cells, progenitor cells of more differentiated Megakaryocytes and Erythrocytes.

### Long lived resident cells show a high adaptation rate in adults

In contrast with the Developmental Immune Cell atlas, where we found mostly progenitor populations with elevated adaptation rates, the picture was very different in the Adult Tissue Immune atlas. Here, we found only four cell types with a high adaptation rate, belonging to the most differentiated and long-lived immune populations (Figure 3, Supplementary Figure **??** and Supplementary Tables **??**, **??**). In the T cell lineage (Figure 3B), the cell types with highest adaptation were tissue resident-memory effector memory CD8+ T cells (Trm em CD8) and resident-memory Th1 and Th17 (Trm Th1 Th17). Trm T cells are long-lived cells that are resident at sites of prior antigenic exposure, i.e. epithelial barrier tissues (lung, skin, gut), to support tissue surveillance and as protective immunity to reinfection[50,51]. For example, influenza-specific lung Trm T cells localise in clusters close to lung epithelial cells, responding rapidly to reinfection through the direct release of cytotoxic mediators, cytokines and chemokines[51]. In humans, higher frequencies of CD8+ Trm cells in the airway have been associated with improved viral control and reduced symptom severity. Trm T cells express different marker genes with potential differences in selection during host-pathogen evolutionary pressure[51]. We therefore reasoned that Trm T cell populations with high protein adaptation could point to selective processes specific to the barrier tissue and pathogen entry site. Intriguingly, both Trm cells with a high protein adaptation were enriched in lymphoid tissue and also in non-lymphoid tissue including liver and lungs, in contrast to Trm cells without signal for adaptation (Trm Tgd and Trm gut CD8) that mostly localised in the gut (Supplementary Figure **??**). Noteworthy, the presence of the Trm T cell populations with a high protein adaptation in the liver and lungs had been validated in situ through targeted spatial profiling[28]. These results suggest a high adaptation rate related to a tissue-specific cellular defence against respiratory pathogens, to provide a rapid response against airborne viruses and bacteria, thereby limiting their growth and spread through the organism.

**Figure 3.**
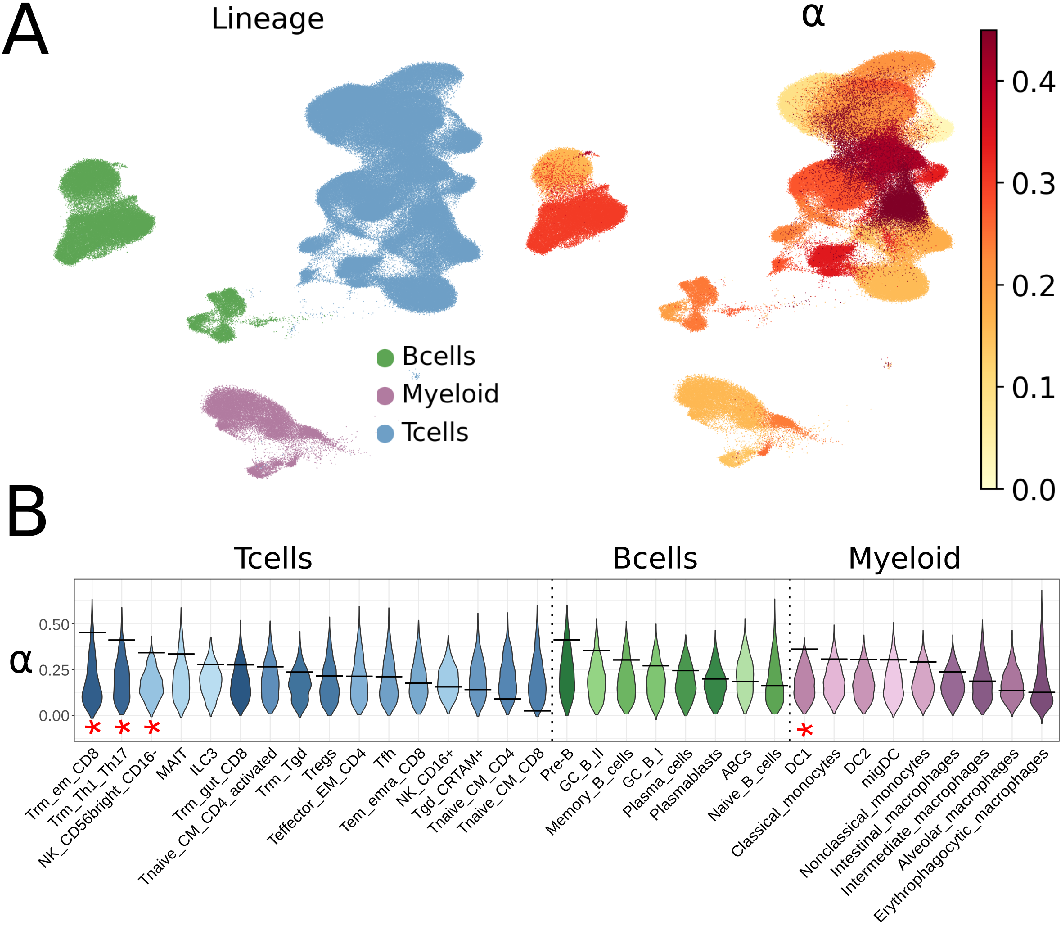
Protein adaptation in the Adult Atlas. A) On the left, each compartment is highlighted in the global UMAP (from Domínguez Conde et al.[28]). On the right, the inferred *α* value is projected on the same UMAP. Note that the *α* value is inferred at the cell-type level. B) For each cell type the inferred *α* value is shown as a horizontal bar and the *α* value distribution of its control bootstrap dataset (*n* = 1000) is shown as a violin plot. The asterisks highlight the cell types with a significant high *α* (* p-value *<* 0.05).

Consistently, the other cell types showing a high adaptation rate in adults are also found in barrier tissues. CD56-bright CD16-negative Natural Killer cells (NK CD56bright CD16-; NK CD16neg) also show tissue-resident profiles and are specialised in cytokine production, in contrast to NK CD16pos cells with cytotoxic activity mostly found in peripheral blood[52]. The role of NK CD16neg cells has been described as regulatory, influencing both innate immunity through cytokine production, but also contributing to the local adaptive immune response[52]. Dendritic cells (DCs) are also found in barrier tissue, such as the lung mucosa, where they sample antigen for their presentation to adaptive immune cells in tissue-draining lymph nodes. Together, we found a high adaptation rate in adult immune cells mainly in cell types localised to barrier tissue as a first line of defence against pathogens, leading us to speculate that the main selective pressure in adult humans comes from external challenges, potentially respiratory pathogens entering the lung epithelium, and that host adaptation is most efficient when it affects early stages of infection before pathogens spread broadly.

### Pathogen-associated early response in Macrophages show a high adaptation rate

To further delineate the type of the selective pressure and the immune response timing leading to adaptation in human immune cells, we charted the dynamics to different stimuli that mimic different pathogen exposure. To this end, we retrieved data from the Human Induced Pluripotent Stem Cells Initiative (HipSci), a project generating human induced pluripotent stem (iPS) cells from from hundreds of individuals, as a resource for directed cell type differentiation and targeted perturbation to chart disease mechanisms[53]. More specifically, we used data from a study involving *>* 200 iPSC lines differentiated into Macrophages and exposed to 10 stimuli that emulate responses to pro/anti-inflammatory cytokines and bacterial or viral infections at two different timepoints (Figure 4A-B). This data allowed us to test the protein adaptation rate of different Macrophages activation states, one of the most plastic immune cell types[54]. To identify processes and timing under adaptation, we evaluated the DEG from Macrophages stimulated for 6h and 24h vs control unstimulated samples. For each stimulus, we categorised responses as “early” (exclusively after 6h), “late” (exclusively after 24h) or “sustained” (found at both time points)[55,56]. This classification simulates the sequential release of inflammatory mediators by Macrophages in response to pathogen products, which constitutes a crucial part of the innate response against invading pathogens. It is worth noting that while an initial inflammatory response is vital, restoring macrophages to their resting state is equally important, as prolonged Macrophage activation leads to tissue damage[57]. Our results (Figure 4C, Supplementary Figure **??**, Supplementary Table **??**, **??**) show high protein adaptation mainly in the early responses, specifically to stimulation with IFNG, R484 and CIL. IFNG is known to induce a pro-inflammatory response in Macrophages associated with intracellular pathogens, including bacterial, protozoal or viral infections. R848 is an agonist of Toll-like receptor (TLR) 7 and 8 that mimics a viral infection, whereas CIL represents a combination of stimuli (CD40 ligand + IFNG + sLPS) to mimic response to bacterial infection. In addition to adaptation signals in the early responses, we found the sustained response to Interleukin 4 (IL4) to have a high adaptation rate. IL4 is generally viewed as a T helper 2 cytokine capable of polarising macrophages into an anti-inflammatory phenotype[58].

**Figure 4.**
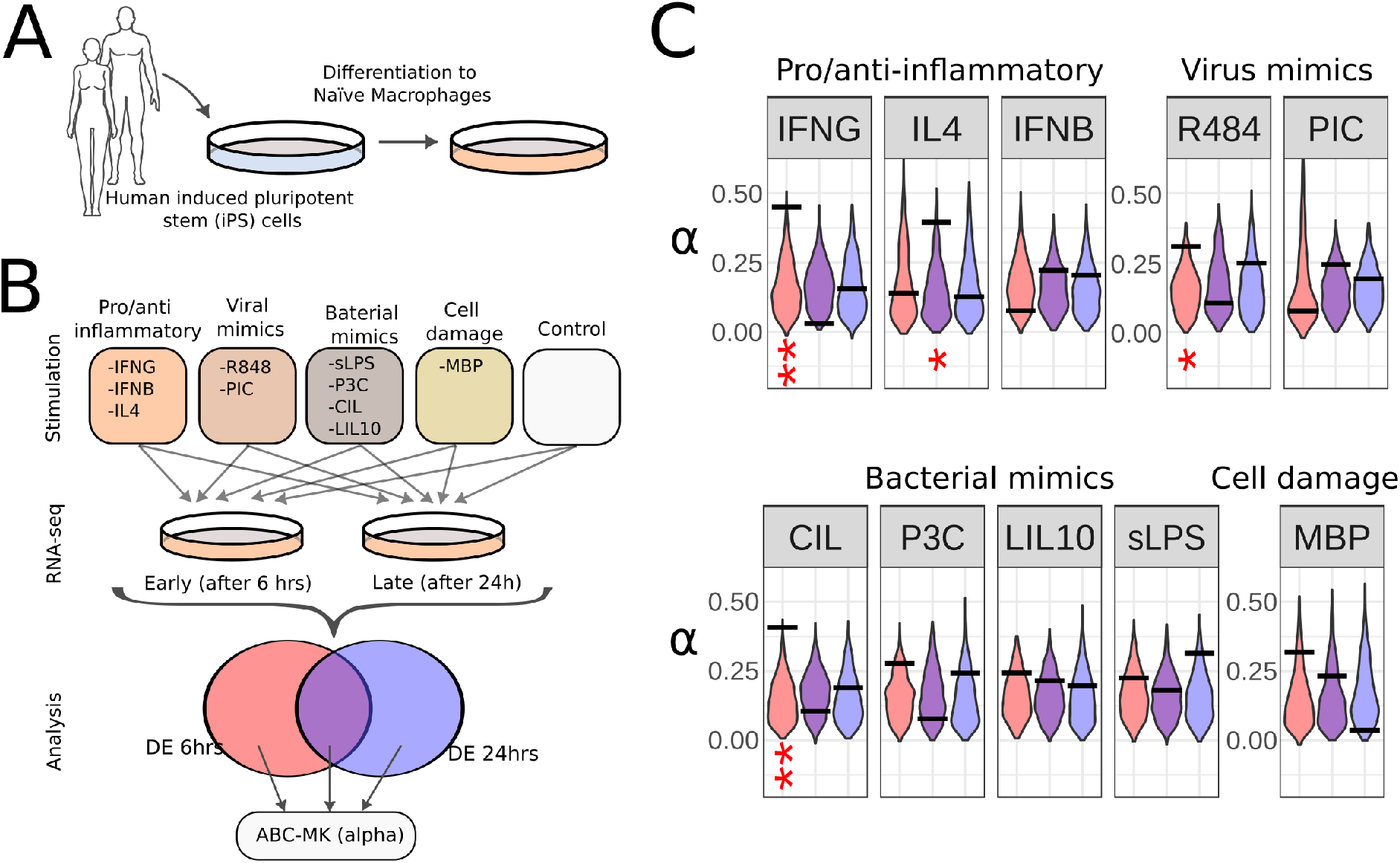
Protein adaptation in Activated Macrophages. A) Overview of the experimental workflow followed by Panousis et al. [54]. More than 200 iPSC lines were derived from unrelated healthy donors and differentiated into naive macrophages. B) Naive macrophages were stimulated and their transcriptomes were sequenced by RNA-seq at two post-stimulation time points (6h and 24h). A differential expression analysis was performed between each timepoint/stimuli and a control. For each stimulus, we further categorised responses as “early” (exclusively after 6h), “late” (exclusively after 24h) or “sustained” (found at both time points). C) For each stimulus and time-specific response we show the inferred *α* value (horizontal bar) and the distribution of the control bootstrap dataset (n=1000) as a violin plot. The asterisks highlight the responses with a significant high *α* (* p-value *<* 0.05, ** p-value *<* 0.01).

### Immune cells are enriched with Eutelostome genes

Having charted the immune types with significant adaptation rates, we then focused on the evolutionary history of genes expressed in the human immune cells. Previous studies[59,60] have shown that evolutionarily younger genes have a higher adaptation rate than older genes. Thus, we reasoned that cell types with high adaptation to be similarly enriched and expressing evolutionary younger genes. To test this hypothesis, we used a phylostratigraphy approach[25], where genes are ranked in different categories or phylostrata (PS) based on their evolutionary emergence time. For humans, nineteen phylostrata have been defined: the first phylostratum (ps1) containing the oldest genes shared by all living organisms, while the last phylostratum (ps19) harbours genes that appeared in the primate lineage and are shared by humans, chimpanzees and marmosets, among others. Using PS value estimates for all human genes[61], we tested for PS enrichment in DEGs of each immune cell type using a hypergeometric test. Our results showed that most immune cell types, in both developmental and adult atlases, are enriched with genes of the Euteleostome phylostrata (PS 12, Figure 5). Euteleostomes (sometimes referred as “bony vertebrates”) is a clade that emerged *>* 400 Mya, which includes jawed fish and their descendants: bony fish, amphibians, reptilians, birds and mammals. Interestingly, it is believed that the adaptive immune system originated in the first jawed fish (the placoderm) as jawed fish descendants have T cell receptors (TCR), B cell receptors (BCR) as well as Major histocompatibility complex (MHC) genes as a consequence of major macroevolutionary events: two whole genome duplications and the invasion of the RAG transposon[62]. Only a few innate immune cell types showed enrichment of younger PS. Macrophages MHCII high, Promonocytes (progenitors), GMP (progenitors), Type 1 innate T cells, and NK cells were enriched with Amniote genes (PS 14) and NK cells was the only cell type enriched with PS 17 genes (subgroup of placental vertebrates that include hoofed and pawed animals).

**Figure 5.**
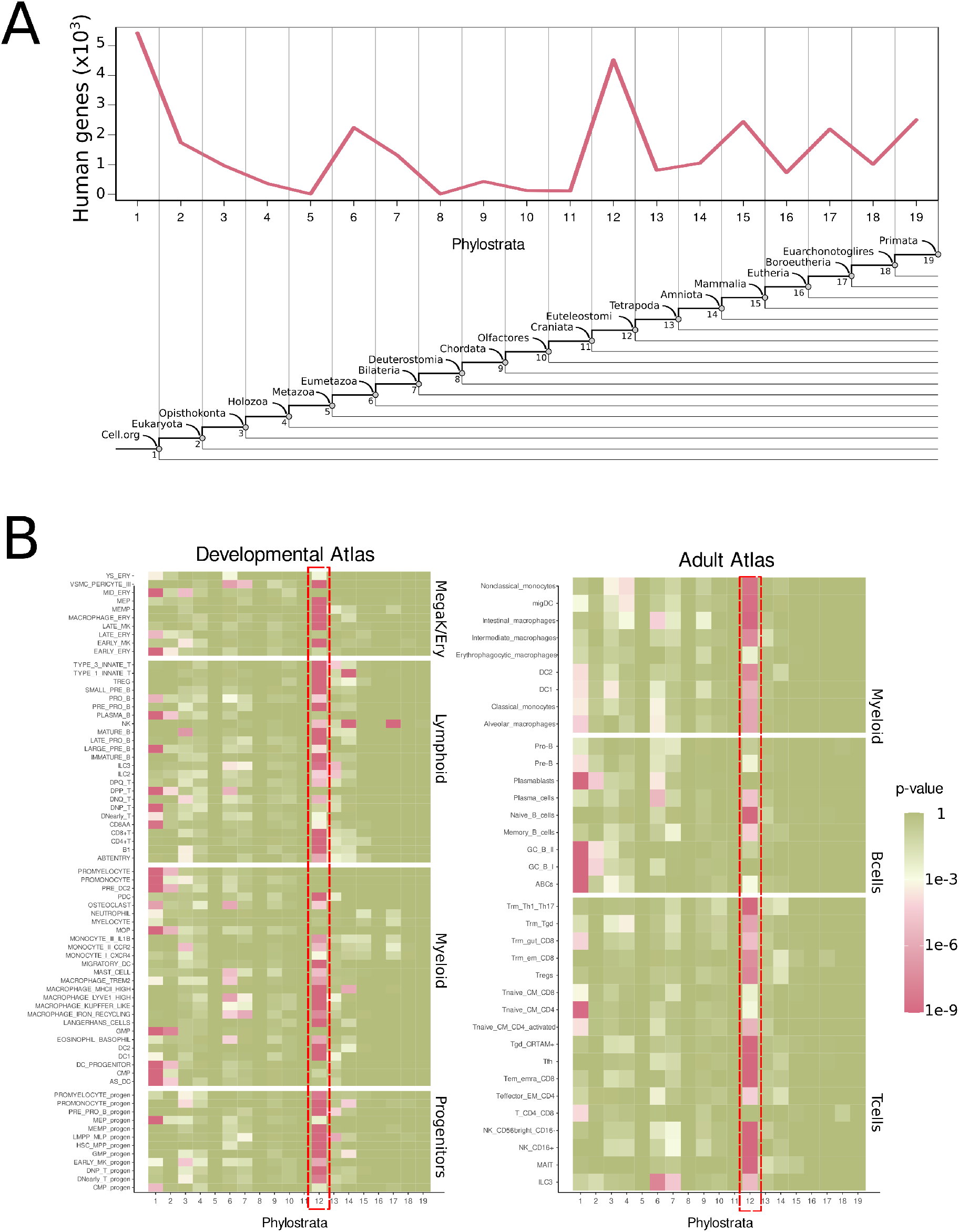
Phylostrata enrichment in immune cell types. A) The line plot shows the number of human genes ranked by their phylostratum (evolutionary time of origin) shown on the x-axis (PS values from[61]. At the bottom, a phylogenetic tree is shown with the clades that correspond to each phylostratum (PS1-PS19). B) Heatmaps showing PS enrichment in the top differentially expressed genes of each immune cell type (hypergeometric test, p-value *<* 0.001) in the Developmental (left) and Adult (right) atlases. The red dotted rectangles highlight PS 12 (Eutelostomes), enriched in most immune cell types.

## Discussion

We combined spatio-temporal transcriptomic dataset of the human immune system and state-of-the-art methods to infer adaptation at the protein sequence level. Looking at the developing and mature immune system through the lens of adaptation provided intriguing insights into the selective pressures that shaped our immune system, including the evolutionary arms race between the human host and its pathogens. Diverging from prior research, we discerned not genes, but cells exhibiting an adaptive signature. Generating an Atlas of Human Adaptation at the cell level, enabled to pinpoint cell types and states under selective pressure and forces that shaped the human innate and adaptive defence since humans split from chimpanzees.

Although the main function of the immune system is defence against pathogens (e.g viral and bacterial), immune cells also play a crucial role in other processes, such as homeostasis and development, potential additional targets of selection. Such functions are orchestrated by highly specialised states at different time points and locations during human development and throughout our lifetime. Given the wide phenotypic and functional diversity of human immune cells with their unique importance towards forming a universal defence mechanism, one might have expected evolutionary pressure to be ubiquitous. However, we found the highest adaptation signals at the very extreme ends of lineage commitment: in early foetal progenitors and in adult tissue-resident cells that form immune memory and represent the first line of defence against repeated infections. These results suggest that selective pressure is strongest at the foundation and executive roles, i.e. cells ensuring immune cell renewal and homeostasis, and at the battle front (barrier tissues), where rapid responses against pathogens are needed.

During development, the foetus is inside an immune-privilege environment, protected by the placenta that serves as an immunological barrier between the foetus and the mother. The immune system is still relatively immature in newborns and matures through childhood until reaching a mature state in adults. The foetal environment is not sterile however and foetal immune cells are exposed to a diverse range of immune-stimulatory molecules, from semi-allogeneic antigens from maternal cells to bacteria. Intriguingly, although stem and progenitor cells are present throughout our lifetime, we only detected high adaptation rates in the foetal population, further underlining the value of the spatio-temporal design and the respective efforts to generate atlases of the developing and adult human body. It further highlights the fact that challenges and tasks of immune cells change continuously during our life to tackle intrinsic and external challenges[63]. In line with this, the response to stimuli in the foetus can be different from the adult. For example, adult DCs respond to toll-like receptor stimulation inducing effector T cells, whereas foetal DCs induce Treg cells as a mechanism of tolerance and immune suppression[64].

We detected genes expressed in foetal HSCs and most HPCs to have a high protein adaptation rate, a surprising result considering their major role to fuel lymphoid and myeloid lineages. HSCs, however, also function as an integrating sensory hub of the immune system, responding to environmental cues within their niche, which is highly distinct in foetal and adult stages[65]. Foetal HSCs arise in different organs at different developmental stages, including the yolk sac, placenta and foetal liver, before colonising the bone marrow in postnatal stages[66]. It has been suggested that these dramatic changes in local environments during the foetal period influence HSC activity[66]. Thus, the requirement of the HSCs and lineage progenitors to respond to a changing environment and challenges in a strictly controlled manner is a plausible candidate as a trait under selection. Additionally, early developmental events can determine later effector responses, as shown by a study in mice, where foetal immune cells responded to prenatal inflammation by activating a transient lymphoid-biased HSC commitment that reshaped not only the foetal, but also postnatal immune output[67]. This opens the possibility that early differentiation events in progenitor cells might be under selection if these regulate the amount of specific immune cell types, especially when the demand for specific amounts of immune cell types has changed during primate evolution—the relative proportions of immune cell types found in homeostasis in adults significantly differ in humans compared to non-human primates[68].

At the other end of immune cell type differentiation, we found adult long-lived tissue resident memory (Trm) T cells with a high rate of protein adaptation. Trm T cells localise in barrier tissues (sites of recurrent infections) and leverage their immune memory from initial encounters with pathogen-derived antigens to provide rapid response to future reinfections. The role of Trm T cells is tightly linked to IFNG signalling. In mice, IFNG produced by Trm cells has been shown to enhance leukocyte recruitment to the infection site and tissue-wide antiviral responses like the upregulation of type I IFN signalling pathway factors[51]. Airway CD8 Trm cells produce IFNG faster than systemic effector CD8 T cells and IFNG-deficient CD8 Trm cells were less effective at controlling pathogen load after viral infection[69]. Similarly, natural Killer CD16neg cells (another lymphoid resident cell type with high adaptation rate) also produce a high amount of IFNG when exposed to bacterial infections[52]. IFNG produced by T and NK cells in turn activates Macrophages, which secrete a high level of inflammatory cytokines and further promote an inflammatory innate response. IFNG activated Macrophages, also known as “classically activated Macrophages”, have a well documented role in host defence to intracellular pathogens including bacterial, protozoal or viral infections (reviewed in Mosser and Edwards[70]).

The high adaptation of resident memory IFNG producing cells, together with our observation that the highest adaptation in activated Macrophages was related to the early response to IFNG, suggests that tissue resident cell types responding to pathogens via IFNG signalling have been under strong adaptive evolution in the human lineage in the millions of years, since humans split from chimpanzees. Based on these results, we hypothesise that host immune adaptation is most efficient in cells residing in recurrent infection sites, to facilitate a prompt immune response before invading pathogens spread, mutate and counter-adapt.

Finally, our phylostratigraphy results suggest that the considerable protein adaptation found in various human immune cell types not to be a consequence of the recruitment of new genes (which tend to evolve and adapt faster), but rather due to adaptive mutations in immune genes that have been present in our ancestors since the emergence of the Immune Adaptive System, more than 400 million years ago.

## Methods

### Gene expression data

scRNAseq annotated data was downloaded in the AnnData format from https://developmental.cellatlas.io/fetal-immune for the Developmental Atlas and from https://www.tissueimmunecellatlas.org/ for the Adult atlas. Specifically, for the developmental Atlas, available H5AD files for the four main immune compartments (HSC/progenitor cells, Megakaryocyte/Erythroid cells, Myeloid cells and Lymphoid cells) were downloaded and used for downstream analysis. For the adult Atlas, available H5AD files for the three main immune compartments (Myeloid, T and Innate lymphoid cells and B cells) were downloaded for downstream differential expression (DE) analyses. The global datasets for both atlases were also downloaded as H5AD files only to produce the UMAP plots in Figure 2A and Figure 2A.

### scRNAseq pre-processing and DE analysis

For each compartment analysed, we discarded cells originally annotated as proliferating or “cycling” (as a cell in a proliferating state expresses a battery of genes related to the cell division process that are not cell-type specific), doublets or low quality. We performed normalisation and log-transformation of the raw data using the scanpy.pp.normalize per cell() and scanpy.pp.log1p() functions of the scanpy package (version 1.9.1;[71]). For differential expression of the different cell types within a compartment, we ran a DE test using the scanpy.tl.rank genes groups() function of the scanpy package, using the wilcoxon test with a p-value threshold of 1 *·* 10*^−^*^4^ and a minimum log2fc value of 1. We then sorted the DE genes by their log2fold change values.

We exported the top 500 DE genes of each cell type (or all genes in case there were fewer than 500) to be analysed with the ABC-MK software. For the developmental atlas we converted the gene ID to ENSEMBL IDs using the ensembldb Bioconductor package (version 2.18.1,[72]). This was not necessary for the Adult atlas, as the genes were already annotated using ENSEMBL IDs.

### Activated macrophages data

Macrophage activation differentially expressed genes were obtained from the study by Panousis et al.[54], available as Supplementary material. As described in the study, iPSCs were differentiated to macrophages[73] and stimulated for 6h and 24h with 10 ng/mL recombinant human interleukin-10 (Peprotech, 200-10-2), 10 ng/mL recombinant human interferon-b (Peprotech, 300-02BC-5), 20 ng/mL recombinant human interleukin-4 (Peprotech, 200-04-5), 50 ng/mL P3C (Pam3CSK4) (Tocris, 4633/1), 20 ng/mL recombinant human interferon-*gamma* (Peprotech, 300-02-20), 10 ng/mL lipopolysaccharides from Escherichia coli O127:B8 (Sigma Aldrich, L3129), 40 ng/mL human recombinant tumour necrosis factor alpha (Peprotech, 300-01A-10), 100 ng/mL R848 (Resiquimod) (Invivogen, tlrl-r848), or 5 ng/mL recombinant human sCD40 Ligand (Peprotech, 310-02-10), 27.5 ug/ml poly I:C (Invivogen, tlrl-pic, diluted in Lipofectamine -Thermo Fisher, L3000001- and Opti-MEM media -Thermo Fisher, 31985062-) as described in[54]. RNA libraries were obtained using the low-input bulk RNA-seq preparation for control and stimulated macrophage samples[54]. Differentially expressed genes for each condition were obtained using the Wald test from DESeq2[74]. Benjamini-Hochberg procedure was used for multiple testing correction (FDR).

From these DE gene lists, we only considered genes with a log2fc *>* 1 and with a p-value *<* 0.05. For each stimulus, we further categorised the genes as “early” (found exclusively in DE gene list after 6h), “late” (found exclusively in DE gene list after 24h) or “sustained” (found at both time points).

### Phylostratigraphy enrichment analysis

We obtained the Phylostrata (PS) values for human genes from the study of[61], where the authors determine consensus ages for human protein-coding and noncoding genes, using publicly available databases. The PS data was downloaded from the supplementary material provided with their original paper.

To test for PS enrichment, we used the hypergeometric test considering all the genes with PS values as the population (*N* = 27, 974) and the DE genes of each cell type (obtained as described above) as the sample (*n ≤* 500). For each PS (1-19) this test uses the hyper-geometric distribution to determine the statistical significance of having drawn a sample of *k* genes of a given PS (out of *n* genes of the sample), from a gene population of size *N* containing *K* number of PS genes. We considered significant p-values *≤* 1 *·* 10*^−^*^3^. The test was performed using the phyper() function of the R stats package (R version 4.1.0).

### Divergence and polymorphic data

Classical and derived MK-test approaches combine both divergence (D) and polymorphism (P) data comparing alleles that are likely to have fitness effects (putatively selected; N) to those less likely to be under selection (putatively neutral, S). To perform MK-test estimations on human immune cell lines, we obtained associated information on polymorphism and divergence data treating synonymous and nonsynonymous mutations as putatively neutral and selected respectively in the protein-coding sequences of the human hg38 assembly and Ensembl release 109 annotations[75].

Divergence counts were obtained by estimating fixed substitution in the human branch based on human, chimpanzee, gorilla, and orangutan alignments. Transcript sequences were obtained by aligning human CDS transcripts from Ensembl v109 on panTro6, gorGor6, and ponAbe3 assemblies from UCSC using pblat[76]. Best reciprocal orthologs were realigned using MACSE v2[77], an explicit multiple sequence alignment software accounting for the underlying codon structure, frameshift and stop codons. The number of non-synonymous and synonymous fixations on the human branch was estimated using Hyphy v2.5[78].

1000GP phase 3 high-coverage phased data[35] across seven African ancestry populations were used to retrieve polymorphic sites and derived allele frequencies. We annotated nonsynonymous and synonymous polymorphism using VEP[75] and estimated ancestral and derived allele frequencies using Ensembl v109 human ancestral allele information from EPO multi-alignments. In total, 17,538 orthologues were included in the analysis, counting sites in multiple isoforms only once.

Because we are interested in the long-term adaptation of the human immune system, we filtered the differentially expressed genes using mammalian orthologs. Gene age can significantly impact the rate of adaptation in populations since populations further away from their optimal conditions take larger evolutionary steps, and those closer to their optimum conditions take smaller steps[60]. This effect is known as the adaptive walk model of gene evolution, exemplified by[60] using Drosophila melanogaster and Arabidopsis thaliana polymorphic and divergence empirical data. We filter differentially expressed genes with the ortholog gene list described in[34] to test for long-term adaptation while avoiding confounding signals due to gene age. This list contains Ensembl v99 human genes with orthologs in a minimum of 251 out of 261 mammalian genome assemblies. As exposed in Di et al.[34], the assemblies were extracted from NCBI Genome and had a minimum N50 contig size of 30kb to reduce the number of possible truncated genes. The orthologs for mammals were defined based on the best-reciprocal hits using Blat[79] alignments, resulting in 13,495 orthologs.

### Quantifying adaptation using ABC-MK

We measured the long-term rate of protein adaptation using ABC-MK, an Approximate Bayesian Computation extension of the MK-test[22,23,34]. Such a rate is usually estimated as the proportion of adaptive non-synonymous substitution and can be derived from classical MK-test by the quantity *α* [14]

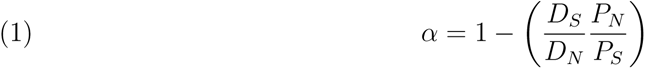

where *D_S_* and *D_N_* are synonymous and non-synonymous fixed differences, and *P_S_* and *P_N_*are synonymous and non-synonymous polymorphic sites. When *α* is close to 1, positive selection is the predominant determinant of molecular divergence. If *α* is close to 0, drift dominates sequence divergence. Nonetheless, *α* approximation from MK-test has multiple drawbacks that could bias the estimation. During the last decade, empirical and theoretical studies have mainly argued that the presence selected alleles segregating can attain at both low and high frequencies leading to an underestimation of *α* [22,80–84]. ABC-MK use a generic Approximate Bayesian Computation (ABC) to infer the rate of adaptation: first, samples the parameter values from prior distributions; second, simulates a model using these random parameter values; and third, calculates informative summary statistics and compares them to observed empirical data. ABC-MK procedure computes the analytical expectation of *α* by frequency categories (*x*) on the Site Frequency Spectrum (similar to[24]), quickly estimating *α*_(_*_x_*_)_ for several combinations of prior distribution values (see Supplementary Table **??** to check prior values). Parameter combinations allow flexibility on the underlying DFE accounting for different amounts of deleterious, weakly and strongly adaptive mutations and background selection (BGS) to finally match the observed *α*_(_*_x_*_)_ with the best-fitting ones among the analytically estimated *α*_(_*_x_*_)_. It is worth noting that weakly adaptive mutations do not reach fixation quickly. Therefore, the existence of non-synonymous polymorphism in the site frequency spectrum (SFS) cannot be ignored, as it can be in the case of strongly adaptive mutations[22]. Moreover, as shown in Uricchio et al.[22], the strength of BGS can also remove weakly adaptive alleles, so linked selection can heavily affect the *α*_(_*_x_*_)_ and the expected amount of fixations. Both disturb the shape of the *α*_(_*_x_*_)_, which translates into a downward trend of *α*_(_*_x_*_)_ at higher frequencies. Accounting independently for weak and strong adaptation along with BGS, ABC-MK can estimate the proportion of adaptation driven by weak and strong adaptation in any virtual DFE scenario, depending on prior values. Note that the mean strength and shape of the gamma deleterious DFE were set from uniform distributed values while accounting for the inferred values in[85], creating random deleterious DFE shapes and amount of weakly deleterious polymorphism. Here we use these functionalities as previously shown in[22,31], to quantify *α* as the contribution of weak and strong adaptive mutation in the presence of BGS: *α* = *α_W_* + *α_S_*. In addition, we also estimated *ω_a_*, the rate of adaptive non-synonymous substitution relative to the mutation rate, following Galtier[83] to support evidence of protein adaptation. For each analysis we subset 10^5^ values from prior distributions to simulate and estimate *α*_(_*_x_*_)_ summary statistics, which allow flexibility for the underlying DFE and BGS strength. Posterior distributions were estimated using ABCreg[30] comparing ABC-MK summary statistics and the empirical datasets. We set tolerance in ABCreg to record 2,500 acceptances from the rejection method and post-adjusted parameters using local linear regression[30,86]. Similarly to[22], *α*_(_*_x_*_)_ summary statistics from the analytical estimations and empirical data were compared using a subset of used *α*_(_*_x_*_)_ values at derived allele counts (DAC) *x* = 2, 4, 5, 10, 20, 50, 200, 661, avoiding singletons since they are particularly sensitive to sequencing errors and distortions due to demographic processes. In addition, note that we exclude any polymorphic site above frequency *x* 0.7 (DAC *>* 925) from empirical and analytical estimations. Based on the findings of[87] about mispolarization regarding the human genome, it can lead to a significant decline in the *α*_(_*_x_*_)_ at higher frequencies, particularly when considering positive selection.

### Bootstrapping

To highlight evolutionary patterns associated with an increased adaptation rate on differentially expressed genes on immune human cell lines, we compare each dataset with 1000 control datasets while matching the total number of analysed genes for each line. Control datasets represent null distribution, accounting for the same average values of multiple confounding factors that can also determine the rate of adaptive evolution compared to the rest of the genome. The bootstrap test uses the same algorithm described by Di et al.[31], Enard and Petrov[32] to build the control sets. The procedure involves adding control genes by pairs to the control set as it grows, ensuring that it has the same range of confounding factor values as the gene set being tested. This approach avoids matching set genes individually with control genes, which would limit the potential gene pool. Instead, we progressively match the gene set with pair controls and set a 5% matching limit over the gene set average confounding values. Additionally, we restrict each control gene from appearing more than three times in each control set as we increase the control pool. We matched a total of 14 confounding factors between set and control genes:

- Ensembl v99 longest CDS length for the gene (between isoforms).
- Average GC content for each coding sequence.
- Average GC1 content at the first codon nucleotide position for each coding sequence.
- Average GC2 content at the second codon nucleotide position for each coding sequence.
- Average GC3 content at the third codon nucleotide position for each coding sequence.
- Average GTEx v8 TPM (Transcripts Per Million) mRNA expression across 53 tissues (in log base 2)[88].
- Average expression (in log base 2) in testis from GTEx v8 tissues THE GTEX CONSORTIUM[88]
- Number of protein-protein interactions (PPIs) in the human protein interaction network[89]. We use the log (base 2) of the number of PPIs.
- Proportion of GERP conserved segments in a 50kb window centered on the Ensembl gene (half way between gene start and gene end)[90].
- Proportion of GERP conserved segments in a 500kb window centered on the Ensembl gene (half way between gene start and gene end)[90]
- Average GERP conservation score in a 50kb window centered on the Ensembl gene[90]
- Average GERP conservation score in a 500kb window centered on the Ensembl gene[90]
- McVicker’s B estimate of background selection in a 50kb window centered on the Ensembl gene[91].
- Average recombination rate in a 500kb window centered on the Ensembl gene, from the deCode 2019 recombination map[92].

## Data availability

We provide the scripts necessary to replicate our analysis in the following Github repository: https://github.com/irepansalvador/Immune Adaptation Atlas 2023.git.

## Acknowledgements

We would like to thank the members of the Single Cell Genomics lab in the CNAG and Antonio Barbadilla for useful comments and feedback. Also we thank Marta Coronado-Zamora and Aina Vaquer for suggesting using the consensus PS dataset.

