## Supplementary Material for "Quantifying Adaptive Evolution of the Human Immune Cell Landscape"

1

### Supplementary materials

**Supplementary Table 1.** Inferred $\alpha$ value and 95% confidence interval for each cell lineage posterior distribution in the Developmental Atlas.

| Lineage | Cell | $\alpha_W$ | $\alpha_S$ | $\alpha$ | Pvalue $\alpha_W$ | Pvalue $\alpha_S$ | Pvalue $\alpha$ |
|---|---|---|---|---|---|---|---|
| HSC progenitors | CMP | 0.114 [-0.002-0.459] | 0.228 [0.054-0.469] | 0.412 [0.182-0.681] | 0.016 | 0.011 | 0.011 |
| HSC progenitors | DNP T | 0.038 [-0.025-0.273] | 0.055 [-0.016-0.238] | 0.138 [0.013-0.396] | 0.691 | 0.653 | 0.592 |
| HSC progenitors | DNearly T | 0.026 [-0.018-0.266] | 0.048 [-0.015-0.254] | 0.103 [-0.006-0.399] | 0.785 | 0.692 | 0.717 |
| HSC progenitors | EARLY MK | 0.1 [0.008-0.366] | 0.133 [0.044-0.344] | 0.328 [0.15-0.544] | 0.052 | 0.090 | 0.022 |
| HSC progenitors | GMP | 0.134 [0.047-0.427] | 0.133 [0.053-0.342] | 0.333 [0.183-0.609] | 0.012 | 0.158 | 0.049 |
| HSC progenitors | HSC MPP | 0.094 [0.002-0.376] | 0.145 [0.039-0.374] | 0.322 [0.157-0.582] | 0.042 | 0.068 | 0.026 |
| HSC progenitors | LMPP MLP | 0.087 [0.008-0.445] | 0.184 [0.05-0.44] | 0.37 [0.156-0.648] | 0.088 | 0.039 | 0.038 |
| HSC progenitors | MEMP | 0.071 [-0.021-0.413] | 0.177 [0.037-0.445] | 0.356 [0.162-0.613] | 0.218 | 0.039 | 0.022 |
| HSC progenitors | MEP | 0.11 [0.03-0.353] | 0.098 [0.032-0.287] | 0.234 [0.113-0.516] | 0.022 | 0.261 | 0.193 |
| HSC progenitors | PRE PRO B | 0.105 [-0.027-0.457] | 0.281 [0.044-0.466] | 0.414 [0.267-0.622] | 0.041 | 0.004 | 0.001 |
| HSC progenitors | PROMONOCYTE | 0.106 [0.012-0.387] | 0.137 [0.042-0.352] | 0.343 [0.163-0.562] | 0.053 | 0.119 | 0.030 |
| HSC progenitors | PROMYELOCYTE | 0.118 [0.023-0.439] | 0.127 [0.052-0.387] | 0.347 [0.146-0.654] | 0.025 | 0.216 | 0.088 |
| Lymphoid | ABTENTRY | 0.035 [-0.027-0.349] | 0.051 [-0.027-0.359] | 0.123 [-0.013-0.508] | 0.568 | 0.620 | 0.606 |
| Lymphoid | B1 | 0.095 [0.0-0.386] | 0.164 [0.04-0.382] | 0.351 [0.18-0.561] | 0.065 | 0.036 | 0.008 |
| Lymphoid | CD4+T | 0.078 [-0.04-0.353] | 0.112 [-0.029-0.352] | 0.25 [0.065-0.503] | 0.194 | 0.245 | 0.208 |
| Lymphoid | CD8+T | 0.038 [-0.042-0.327] | 0.086 [-0.027-0.342] | 0.179 [0.018-0.459] | 0.644 | 0.367 | 0.407 |
| Lymphoid | CD8AA | 0.098 [0.013-0.378] | 0.13 [0.048-0.371] | 0.282 [0.128-0.564] | 0.086 | 0.131 | 0.117 |
| Lymphoid | DNP T | 0.033 [-0.063-0.33] | 0.124 [-0.031-0.336] | 0.211 [0.06-0.446] | 0.773 | 0.134 | 0.282 |
| Lymphoid | DNQ T | 0.056 [0.004-0.304] | 0.092 [0.017-0.273] | 0.175 [0.062-0.454] | 0.352 | 0.293 | 0.389 |
| Lymphoid | DNearly T | 0.038 [-0.003-0.264] | 0.042 [-0.003-0.227] | 0.102 [0.013-0.392] | 0.629 | 0.743 | 0.721 |
| Lymphoid | DPP T | 0.046 [-0.075-0.375] | 0.241 [0.017-0.439] | 0.311 [0.151-0.53] | 0.548 | 0.005 | 0.040 |
| Lymphoid | DPQ T | 0.042 [-0.025-0.281] | 0.064 [-0.01-0.264] | 0.16 [0.028-0.415] | 0.577 | 0.529 | 0.452 |
| Lymphoid | ILC2 | 0.067 [0.017-0.365] | 0.052 [0.01-0.266] | 0.148 [0.052-0.554] | 0.290 | 0.699 | 0.576 |
| Lymphoid | ILC3 | 0.135 [0.04-0.417] | 0.121 [0.04-0.33] | 0.307 [0.168-0.604] | 0.006 | 0.139 | 0.034 |
| Lymphoid | IMMATURE B | 0.045 [-0.057-0.42] | 0.19 [0.025-0.481] | 0.358 [0.123-0.62] | 0.496 | 0.031 | 0.038 |
| Lymphoid | LARGE PRE B | 0.027 [-0.082-0.325] | 0.136 [-0.024-0.352] | 0.222 [0.071-0.444] | 0.838 | 0.065 | 0.213 |
| Lymphoid | LATE PRO B | 0.031 [-0.069-0.394] | 0.254 [0.026-0.473] | 0.338 [0.174-0.571] | 0.725 | 0.008 | 0.049 |
| Lymphoid | MATURE B | 0.058 [-0.057-0.407] | 0.267 [0.03-0.451] | 0.343 [0.193-0.576] | 0.395 | 0.002 | 0.020 |
| Lymphoid | NK | 0.123 [0.031-0.401] | 0.115 [0.044-0.328] | 0.283 [0.156-0.601] | 0.020 | 0.218 | 0.104 |
| Lymphoid | PLASMA B | 0.017 [-0.018-0.298] | 0.038 [-0.024-0.302] | 0.074 [-0.018-0.449] | 0.845 | 0.739 | 0.810 |
| Lymphoid | PRE PRO B | 0.11 [0.009-0.406] | 0.144 [0.034-0.353] | 0.316 [0.179-0.572] | 0.028 | 0.074 | 0.040 |
| Lymphoid | PRO B | -0.016 [-0.131-0.335] | 0.275 [0.011-0.442] | 0.275 [0.122-0.481] | 1.000 | 0.001 | 0.103 |

| | | | | | | | |
|---|---|---|---|---|---|---|---|
| Lymphoid | SMALL PRE B | 0.081 [-0.023-0.37] | 0.128 [0.015-0.385] | 0.249 [0.11-0.545] | 0.135 | 0.134 | 0.183 |
| Lymphoid | TREG | 0.101 [-0.006-0.434] | 0.239 [0.04-0.432] | 0.378 [0.209-0.617] | 0.063 | 0.012 | 0.015 |
| Lymphoid | TYPE 1 INNATE T | 0.111 [0.019-0.47] | 0.213 [0.06-0.485] | 0.39 [0.225-0.694] | 0.038 | 0.033 | 0.039 |
| Lymphoid | TYPE 3 INNATE T | 0.069 [-0.029-0.398] | 0.146 [0.001-0.403] | 0.293 [0.109-0.551] | 0.281 | 0.108 | 0.117 |
| MegaK Ery | EARLY ERY | 0.124 [0.041-0.391] | 0.116 [0.05-0.323] | 0.298 [0.154-0.571] | 0.010 | 0.183 | 0.079 |
| MegaK Ery | EARLY MK MegaK Ery | 0.064 [-0.031-0.372] | 0.181 [0.008-0.365] | 0.307 [0.154-0.506] | 0.315 | 0.023 | 0.039 |
| MegaK Ery | LATE ERY | 0.057 [-0.01-0.41] | 0.13 [0.019-0.466] | 0.302 [0.072-0.628] | 0.248 | 0.135 | 0.120 |
| MegaK Ery | LATE MK | 0.046 [-0.034-0.315] | 0.093 [-0.02-0.282] | 0.17 [0.036-0.432] | 0.553 | 0.344 | 0.454 |
| MegaK Ery | MACROPHAGE ERY | 0.068 [-0.009-0.362] | 0.109 [0.019-0.346] | 0.212 [0.075-0.536] | 0.363 | 0.340 | 0.415 |
| MegaK Ery | MEMP MegaK Ery | 0.097 [-0.001-0.391] | 0.149 [0.016-0.349] | 0.3 [0.143-0.542] | 0.052 | 0.075 | 0.074 |
| MegaK Ery | MEP MegaK Ery | 0.109 [0.023-0.404] | 0.144 [0.033-0.385] | 0.346 [0.185-0.576] | 0.024 | 0.077 | 0.017 |
| MegaK Ery | MID ERY | 0.04 [-0.045-0.291] | 0.082 [-0.024-0.3] | 0.157 [0.03-0.428] | 0.672 | 0.443 | 0.521 |
| MegaK Ery | VSMC PERI-CYTE III | 0.046 [-0.017-0.267] | 0.065 [-0.002-0.231] | 0.156 [0.04-0.373] | 0.619 | 0.581 | 0.548 |
| MegaK Ery | YS ERY | 0.091 [0.013-0.466] | 0.126 [0.033-0.419] | 0.278 [0.103-0.68] | 0.059 | 0.168 | 0.188 |
| Myeloid | AS DC | 0.025 [-0.071-0.316] | 0.11 [-0.029-0.34] | 0.198 [0.035-0.427] | 0.842 | 0.222 | 0.335 |
| Myeloid | CMP | 0.077 [-0.036-0.372] | 0.156 [0.003-0.354] | 0.28 [0.125-0.492] | 0.158 | 0.076 | 0.121 |
| Myeloid | DC1 | 0.048 [-0.052-0.353] | 0.137 [-0.021-0.355] | 0.263 [0.079-0.476] | 0.471 | 0.078 | 0.103 |
| Myeloid | DC2 | 0.083 [-0.015-0.381] | 0.121 [0.017-0.364] | 0.276 [0.115-0.543] | 0.122 | 0.144 | 0.110 |
| Myeloid | DC PROGENI-TOR | 0.065 [-0.04-0.374] | 0.199 [-0.006-0.398] | 0.286 [0.124-0.542] | 0.275 | 0.023 | 0.083 |
| Myeloid | EOSINOPHIL BA-SOPHIL | 0.079 [0.013-0.321] | 0.081 [0.021-0.261] | 0.194 [0.08-0.471] | 0.144 | 0.382 | 0.318 |
| Myeloid | GMP | 0.056 [-0.011-0.283] | 0.073 [-0.002-0.259] | 0.158 [0.04-0.415] | 0.487 | 0.543 | 0.556 |
| Myeloid | LANGERHANS CELLS | 0.004 [-0.017-0.337] | 0.011 [-0.021-0.324] | 0.031 [-0.031-0.523] | 0.928 | 0.930 | 0.934 |
| Myeloid | MACROPHAGE IRON RECY-CLING | 0.084 [0.03-0.325] | 0.086 [0.029-0.245] | 0.212 [0.097-0.487] | 0.101 | 0.347 | 0.249 |
| Myeloid | MACROPHAGE KUPFFER LIKE | 0.074 [-0.007-0.325] | 0.118 [0.015-0.296] | 0.24 [0.104-0.459] | 0.314 | 0.200 | 0.214 |
| Myeloid | MACROPHAGE LYVE1 HIGH | 0.124 [0.0-0.419] | 0.224 [0.053-0.406] | 0.365 [0.221-0.616] | 0.013 | 0.018 | 0.011 |

| | | | | | | | |
|---|---|---|---|---|---|---|---|
| Myeloid | MACROPHAGE MHCII HIGH | 0.038 [-0.027-0.329] | 0.045 [-0.017-0.297] | 0.141 [-0.008-0.478] | 0.528 | 0.691 | 0.531 |
| Myeloid | MACROPHAGE TREM2 | 0.125 [0.029-0.409] | 0.154 [0.057-0.36] | 0.332 [0.191-0.59] | 0.026 | 0.101 | 0.063 |
| Myeloid | MAST CELL | 0.121 [0.023-0.436] | 0.184 [0.055-0.396] | 0.367 [0.228-0.608] | 0.003 | 0.015 | 0.003 |
| Myeloid | MIGRATORY DC | 0.084 [-0.028-0.347] | 0.126 [0.011-0.336] | 0.254 [0.108-0.505] | 0.177 | 0.164 | 0.190 |
| Myeloid | MONOCYTE III IL1B | 0.103 [0.02-0.397] | 0.139 [0.037-0.378] | 0.325 [0.141-0.583] | 0.053 | 0.128 | 0.068 |
| Myeloid | MONOCYTE II CCR2 | 0.044 [-0.012-0.301] | 0.081 [0.0-0.31] | 0.169 [0.029-0.465] | 0.544 | 0.446 | 0.503 |
| Myeloid | MONOCYTE I CXCR4 | 0.056 [-0.014-0.321] | 0.074 [-0.001-0.302] | 0.188 [0.049-0.458] | 0.366 | 0.478 | 0.372 |
| Myeloid | MOP | 0.048 [-0.035-0.338] | 0.108 [-0.015-0.32] | 0.203 [0.068-0.466] | 0.547 | 0.196 | 0.278 |
| Myeloid | MYELOCYTE | 0.045 [-0.017-0.457] | 0.123 [0.009-0.546] | 0.314 [0.046-0.721] | 1.000 | 1.000 | 1.000 |
| Myeloid | NEUTROPHIL | 0.087 [-0.012-0.486] | 0.224 [0.046-0.536] | 0.393 [0.151-0.726] | 0.050 | 0.037 | 0.061 |
| Myeloid | OSTEOCLAST | 0.033 [-0.005-0.273] | 0.034 [0.0-0.205] | 0.085 [0.013-0.397] | 0.724 | 0.825 | 0.804 |
| Myeloid | PDC | 0.077 [-0.006-0.345] | 0.107 [0.028-0.317] | 0.242 [0.11-0.517] | 0.133 | 0.210 | 0.144 |
| Myeloid | PRE DC2 | 0.045 [-0.058-0.353] | 0.156 [0.001-0.38] | 0.275 [0.118-0.496] | 0.572 | 0.050 | 0.091 |
| Myeloid | PROMONOCYTE | 0.079 [-0.026-0.38] | 0.218 [0.016-0.38] | 0.333 [0.157-0.526] | 0.224 | 0.015 | 0.037 |
| Myeloid | PROMYELOCYTE | 0.046 [-0.013-0.285] | 0.054 [-0.012-0.249] | 0.135 [0.016-0.414] | 0.588 | 0.713 | 0.664 |

**Supplementary Table 2.** Inferred $\alpha$ value and 95% confidence interval for each cell lineage posterior distribution in the Adult Atlas.

| Lineage | Cell | $\alpha_W$ | $\alpha_S$ | $\alpha$ | Pvalue $\alpha_W$ | Pvalue $\alpha_S$ | Pvalue $\alpha$ |
|---|---|---|---|---|---|---|---|
| Bcells | ABCs | 0.077 [-0.008-0.326] | 0.083 [-0.009-0.296] | 0.184 [0.054-0.481] | 0.526 | 0.537 | 0.577 |
| Bcells | GC B I | 0.044 [-0.065-0.376] | 0.15 [-0.028-0.395] | 0.271 [0.096-0.502] | 0.774 | 0.070 | 0.195 |
| Bcells | GC B II | 0.101 [-0.004-0.412] | 0.15 [0.013-0.428] | 0.354 [0.148-0.616] | 0.226 | 0.092 | 0.080 |
| Bcells | Memory B cells | 0.081 [-0.007-0.377] | 0.132 [-0.005-0.371] | 0.302 [0.106-0.533] | 0.336 | 0.167 | 0.160 |
| Bcells | Naive B cells | 0.061 [0.0-0.38] | 0.064 [0.005-0.327] | 0.163 [0.032-0.568] | 0.434 | 0.562 | 0.506 |
| Bcells | Plasma cells | 0.092 [-0.005-0.321] | 0.089 [0.007-0.283] | 0.244 [0.088-0.455] | 0.286 | 0.349 | 0.258 |
| Bcells | Plasmablasts | 0.039 [-0.058-0.301] | 0.09 [-0.033-0.321] | 0.199 [0.036-0.435] | 0.801 | 0.316 | 0.379 |
| Bcells | Pre-B | 0.075 [-0.048-0.461] | 0.181 [0.026-0.563] | 0.411 [0.152-0.688] | 0.480 | 0.094 | 0.096 |
| Bcells | Pro-B | 0.059 [-0.009-0.489] | 0.113 [0.004-0.539] | 0.281 [0.056-0.746] | 1.000 | 1.000 | 1.000 |
| Myeloid | Alveolar macrophages | 0.028 [-0.036-0.263] | 0.062 [-0.021-0.248] | 0.136 [0.013-0.369] | 0.811 | 0.599 | 0.610 |
| Myeloid | Classical monocytes | 0.135 [0.032-0.429] | 0.116 [0.038-0.339] | 0.306 [0.167-0.595] | 0.014 | 0.268 | 0.106 |
| Myeloid | DC1 | 0.041 [-0.084-0.41] | 0.296 [0.028-0.479] | 0.36 [0.198-0.581] | 0.652 | 0.001 | 0.011 |
| Myeloid | DC2 | 0.036 [-0.075-0.367] | 0.22 [0.017-0.422] | 0.305 [0.14-0.523] | 0.733 | 0.014 | 0.075 |
| Myeloid | Erythrophagocytic macrophages | 0.038 [-0.028-0.393] | 0.051 [-0.046-0.431] | 0.126 [-0.034-0.615] | 0.412 | 0.580 | 0.548 |
| Myeloid | Intermediate macrophages | 0.038 [-0.03-0.315] | 0.098 [-0.003-0.328] | 0.186 [0.03-0.47] | 0.659 | 0.312 | 0.424 |
| Myeloid | Intestinal macrophages | 0.071 [-0.019-0.331] | 0.144 [0.033-0.327] | 0.237 [0.119-0.494] | 0.296 | 0.090 | 0.211 |
| Myeloid | Nonclassical monocytes | 0.112 [0.005-0.383] | 0.126 [0.034-0.328] | 0.291 [0.147-0.561] | 0.055 | 0.183 | 0.117 |
| Myeloid | migDC | 0.059 [-0.044-0.378] | 0.146 [0.009-0.426] | 0.303 [0.127-0.562] | 0.419 | 0.128 | 0.126 |
| Tcells | ILC3 | 0.107 [0.044-0.358] | 0.118 [0.046-0.304] | 0.277 [0.15-0.548] | 0.158 | 0.187 | 0.163 |
| Tcells | MAIT | 0.115 [0.02-0.453] | 0.123 [0.027-0.405] | 0.334 [0.124-0.666] | 0.097 | 0.209 | 0.135 |
| Tcells | NK CD16+ | 0.086 [0.029-0.325] | 0.055 [0.013-0.233] | 0.156 [0.069-0.504] | 0.373 | 0.667 | 0.613 |
| Tcells | NK CD56bright CD16- | 0.186 [0.102-0.458] | 0.118 [0.052-0.319] | 0.341 [0.225-0.675] | 0.003 | 0.111 | 0.019 |
| Tcells | T CD4 CD8 | 0.076 [-0.016-0.568] | 0.26 [0.038-0.747] | 0.481 [0.141-0.951] | 1.000 | 1.000 | 1.000 |
| Tcells | Teffector EM CD4 | 0.079 [0.001-0.366] | 0.097 [0.015-0.35] | 0.213 [0.06-0.55] | 0.354 | 0.355 | 0.397 |
| Tcells | Tem emra CD8 | 0.071 [-0.008-0.326] | 0.07 [-0.006-0.284] | 0.176 [0.046-0.462] | 0.431 | 0.531 | 0.501 |
| Tcells | Tfh | 0.062 [-0.013-0.383] | 0.096 [-0.004-0.403] | 0.211 [0.041-0.581] | 0.456 | 0.368 | 0.398 |
| Tcells | Tgd CRTAM+ | 0.057 [0.005-0.305] | 0.07 [0.006-0.265] | 0.141 [0.042-0.476] | 0.610 | 0.579 | 0.661 |
| Tcells | Tnaive CM CD4 | 0.044 [-0.002-0.32] | 0.03 [-0.007-0.24] | 0.09 [0.01-0.474] | 0.744 | 0.881 | 0.849 |

| | | | | | | | |
|---|---|---|---|---|---|---|---|
| Tcells | Tnaive CM CD4 activated | 0.113 [0.034-0.385] | 0.103 [0.015-0.346] | 0.265 [0.129-0.548] | 0.148 | 0.307 | 0.242 |
| Tcells | Tnaive CM CD8 | 0.005 [-0.019-0.268] | 0.007 [-0.032-0.23] | 0.026 [-0.038-0.402] | 0.987 | 0.979 | 0.980 |
| Tcells | Tregs | 0.085 [0.013-0.389] | 0.096 [0.03-0.344] | 0.215 [0.09-0.555] | 0.249 | 0.315 | 0.342 |
| Tcells | Trm Tgd | 0.112 [0.025-0.371] | 0.091 [0.014-0.285] | 0.236 [0.116-0.507] | 0.138 | 0.264 | 0.218 |
| Tcells | Trm Th1 Th17 | 0.189 [0.029-0.53] | 0.159 [0.053-0.453] | 0.41 [0.239-0.739] | 0.003 | 0.096 | 0.044 |
| Tcells | Trm em CD8 | 0.104 [0.021-0.506] | 0.228 [0.054-0.617] | 0.45 [0.225-0.805] | 0.089 | 0.023 | 0.043 |
| Tcells | Trm gut CD8 | 0.105 [0.01-0.39] | 0.126 [0.016-0.352] | 0.276 [0.128-0.559] | 0.176 | 0.133 | 0.183 |

**Supplementary Table 3.** Inferred $\alpha$ value and 95% confidence interval on iPSC lines differentiated into Macrophages and exposed to 10 stimuli that emulate responses to pro/anti-inflammatory cytokines and bacterial or viral infections at two different timepoints.

| Stimuli | Time | $\alpha_W$ | $\alpha_S$ | $\alpha$ | Pvalue $\alpha_W$ | Pvalue $\alpha_S$ | Pvalue $\alpha$ |
|---|---|---|---|---|---|---|---|
| CIL | Early | 0.106 [0.001-0.443] | 0.239 [0.06-0.443] | 0.398 [0.249-0.626] | 0.021 | 0.005 | 0.003 |
| CIL | Inter | 0.044 [-0.02-0.267] | 0.055 [-0.012-0.232] | 0.132 [0.016-0.389] | 0.623 | 0.689 | 0.671 |
| CIL | Late | 0.044 [-0.029-0.306] | 0.082 [-0.007-0.291] | 0.179 [0.038-0.444] | 0.474 | 0.349 | 0.346 |
| IFNB | Early | 0.025 [-0.015-0.285] | 0.034 [-0.011-0.224] | 0.077 [-0.009-0.43] | 0.822 | 0.824 | 0.844 |
| IFNB | Inter | 0.059 [-0.032-0.327] | 0.121 [0.003-0.323] | 0.221 [0.096-0.464] | 0.334 | 0.126 | 0.205 |
| IFNB | Late | 0.042 [-0.048-0.321] | 0.112 [-0.017-0.342] | 0.204 [0.044-0.462] | 0.522 | 0.173 | 0.267 |
| IFNG | Early | 0.151 [0.043-0.487] | 0.18 [0.076-0.454] | 0.45 [0.247-0.708] | 0.002 | 0.041 | 0.003 |
| IFNG | Inter | 0.009 [-0.016-0.269] | 0.009 [-0.024-0.18] | 0.03 [-0.025-0.379] | 0.968 | 0.984 | 0.984 |
| IFNG | Late | 0.036 [-0.046-0.308] | 0.077 [-0.012-0.325] | 0.156 [0.013-0.458] | 0.593 | 0.415 | 0.466 |
| IL4 | Early | 0.025 [-0.025-0.363] | 0.07 [-0.025-0.409] | 0.139 [-0.013-0.563] | 0.615 | 0.441 | 0.496 |
| IL4 | Inter | 0.095 [-0.002-0.452] | 0.226 [0.043-0.484] | 0.395 [0.187-0.667] | 0.031 | 0.013 | 0.021 |
| IL4 | Late | 0.017 [-0.036-0.342] | 0.049 [-0.03-0.348] | 0.126 [-0.033-0.506] | 0.764 | 0.575 | 0.531 |
| LIL10 | Early | 0.058 [-0.046-0.346] | 0.121 [-0.024-0.346] | 0.244 [0.088-0.463] | 0.400 | 0.146 | 0.162 |
| LIL10 | Inter | 0.072 [0.004-0.303] | 0.095 [0.021-0.285] | 0.216 [0.076-0.445] | 0.207 | 0.286 | 0.230 |
| LIL10 | Late | 0.044 [-0.024-0.313] | 0.116 [0.01-0.345] | 0.197 [0.056-0.479] | 0.494 | 0.158 | 0.306 |
| MBP | Early | 0.062 [-0.029-0.441] | 0.176 [0.035-0.487] | 0.319 [0.123-0.652] | 0.182 | 0.047 | 0.086 |
| MBP | Inter | 0.049 [-0.004-0.365] | 0.091 [0.001-0.387] | 0.233 [0.04-0.54] | 0.325 | 0.286 | 0.218 |
| MBP | Late | 0.007 [-0.025-0.278] | 0.017 [-0.021-0.259] | 0.036 [-0.032-0.423] | 0.937 | 0.907 | 0.942 |
| P3C | Early | 0.069 [-0.024-0.364] | 0.156 [0.021-0.373] | 0.278 [0.148-0.527] | 0.225 | 0.056 | 0.079 |
| P3C | Inter | 0.029 [-0.012-0.304] | 0.034 [-0.005-0.23] | 0.078 [0.001-0.449] | 0.695 | 0.787 | 0.798 |
| P3C | Late | 0.017 [-0.091-0.373] | 0.126 [-0.036-0.423] | 0.243 [0.059-0.517] | 0.818 | 0.135 | 0.189 |
| PIC | Early | 0.017 [-0.044-0.424] | 0.023 [-0.064-0.405] | 0.075 [-0.081-0.629] | 0.668 | 0.798 | 0.707 |
| PIC | Inter | 0.066 [-0.013-0.323] | 0.106 [0.015-0.296] | 0.242 [0.088-0.452] | 0.252 | 0.224 | 0.145 |
| PIC | Late | 0.069 [-0.025-0.32] | 0.097 [0.002-0.291] | 0.191 [0.062-0.447] | 0.272 | 0.267 | 0.328 |
| R484 | Early | 0.078 [-0.019-0.383] | 0.161 [0.031-0.391] | 0.308 [0.157-0.558] | 0.110 | 0.038 | 0.040 |
| R484 | Inter | 0.027 [-0.021-0.289] | 0.035 [-0.013-0.253] | 0.103 [-0.014-0.432] | 0.726 | 0.779 | 0.681 |
| R484 | Late | 0.051 [-0.035-0.348] | 0.104 [-0.014-0.359] | 0.248 [0.052-0.497] | 0.402 | 0.237 | 0.187 |
| all | Early | 0.079 [-0.012-0.353] | 0.125 [0.027-0.331] | 0.263 [0.121-0.509] | 0.170 | 0.147 | 0.131 |
| all | Inter | 0.092 [0.025-0.36] | 0.139 [0.042-0.374] | 0.285 [0.138-0.584] | 0.050 | 0.095 | 0.099 |
| all | Late | 0.036 [-0.013-0.242] | 0.051 [-0.011-0.217] | 0.104 [0.01-0.355] | 0.656 | 0.634 | 0.689 |
| sLPS | Early | 0.072 [-0.027-0.328] | 0.107 [0.004-0.312] | 0.226 [0.091-0.461] | 0.243 | 0.238 | 0.225 |
| sLPS | Inter | 0.054 [-0.026-0.299] | 0.083 [-0.003-0.271] | 0.181 [0.05-0.427] | 0.451 | 0.396 | 0.376 |
| sLPS | Late | 0.094 [-0.001-0.385] | 0.143 [0.045-0.378] | 0.314 [0.147-0.552] | 0.060 | 0.081 | 0.059 |

**Supplementary Table 4.** Inferred $\omega_a$ value and 95% confidence interval for each cell lineage posterior distribution in the Developmental Atlas.

| Lineage | Cell | $\omega_{aW}$ | $\omega_{aS}$ | $\omega_a$ | Pvalue $\omega_{aW}$ | Pvalue $\omega_{aS}$ | Pvalue $\omega_a$ |
|---|---|---|---|---|---|---|---|
| HSC progenitors | CMP | 0.019 [-0.025-0.18] | 0.068 [-0.01-0.195] | 0.121 [0.025-0.267] | 0.099 | 0.022 | 0.014 |
| HSC progenitors | DNP T | 0.006 [-0.005-0.07] | 0.01 [-0.005-0.065] | 0.028 [-0.002-0.109] | 0.747 | 0.711 | 0.736 |
| HSC progenitors | DNearly T | 0.003 [-0.005-0.073] | 0.008 [-0.006-0.076] | 0.021 [-0.007-0.122] | 0.958 | 0.807 | 0.879 |
| HSC progenitors | EARLY MK | 0.015 [-0.004-0.105] | 0.026 [0.001-0.107] | 0.064 [0.019-0.162] | 0.142 | 0.165 | 0.127 |
| HSC progenitors | GMP | 0.022 [-0.003-0.137] | 0.031 [-0.003-0.123] | 0.086 [0.017-0.194] | 0.032 | 0.182 | 0.062 |
| HSC progenitors | HSC MPP | 0.022 [-0.012-0.14] | 0.03 [-0.003-0.145] | 0.097 [0.027-0.217] | 0.026 | 0.133 | 0.013 |
| HSC progenitors | LMPP MLP | 0.03 [-0.02-0.194] | 0.063 [-0.007-0.199] | 0.14 [0.032-0.282] | 0.008 | 0.022 | 0.001 |
| HSC progenitors | MEMP | 0.011 [-0.011-0.134] | 0.051 [-0.002-0.156] | 0.096 [0.021-0.201] | 0.316 | 0.035 | 0.032 |
| HSC progenitors | MEP | 0.016 [0.004-0.108] | 0.015 [-0.001-0.088] | 0.054 [0.011-0.156] | 0.115 | 0.499 | 0.288 |
| HSC progenitors | PRE PRO B | 0.026 [-0.013-0.137] | 0.079 [-0.004-0.159] | 0.105 [0.054-0.198] | 0.015 | 0.001 | 0.002 |
| HSC progenitors | PROMONOCYTE | 0.023 [-0.005-0.133] | 0.042 [-0.004-0.121] | 0.081 [0.021-0.184] | 0.027 | 0.075 | 0.077 |
| HSC progenitors | PROMYELOCYTE | 0.03 [-0.021-0.193] | 0.041 [-0.01-0.165] | 0.103 [0.011-0.272] | 0.019 | 0.139 | 0.065 |
| Lymphoid | ABTENTRY | 0.005 [-0.006-0.082] | 0.008 [-0.006-0.082] | 0.028 [-0.008-0.124] | 0.804 | 0.861 | 0.790 |
| Lymphoid | B1 | 0.017 [-0.008-0.113] | 0.05 [-0.004-0.122] | 0.08 [0.021-0.176] | 0.083 | 0.022 | 0.026 |
| Lymphoid | CD4+T | 0.007 [-0.006-0.082] | 0.02 [-0.01-0.093] | 0.047 [0.0-0.125] | 0.621 | 0.383 | 0.461 |
| Lymphoid | CD8+T | 0.003 [-0.006-0.063] | 0.016 [-0.007-0.076] | 0.039 [-0.004-0.104] | 0.983 | 0.473 | 0.573 |
| Lymphoid | CD8AA | 0.016 [-0.005-0.119] | 0.028 [0.0-0.123] | 0.071 [0.01-0.191] | 0.175 | 0.198 | 0.183 |
| Lymphoid | DNP T | 0.005 [-0.01-0.084] | 0.028 [-0.006-0.092] | 0.049 [0.003-0.127] | 0.833 | 0.172 | 0.392 |
| Lymphoid | DNQ T | 0.011 [-0.004-0.112] | 0.02 [-0.001-0.099] | 0.052 [0.004-0.166] | 0.272 | 0.316 | 0.282 |
| Lymphoid | DNearly T | 0.006 [-0.003-0.083] | 0.008 [-0.005-0.075] | 0.027 [-0.004-0.129] | 0.734 | 0.828 | 0.761 |
| Lymphoid | DPP T | 0.01 [-0.014-0.09] | 0.058 [0.003-0.12] | 0.07 [0.027-0.152] | 0.458 | 0.015 | 0.122 |
| Lymphoid | DPQ T | 0.007 [-0.006-0.082] | 0.016 [-0.005-0.086] | 0.045 [-0.001-0.13] | 0.579 | 0.451 | 0.429 |
| Lymphoid | ILC2 | 0.036 [-0.003-0.178] | 0.036 [-0.003-0.138] | 0.074 [0.008-0.289] | 0.012 | 0.166 | 0.207 |
| Lymphoid | ILC3 | 0.024 [-0.012-0.151] | 0.031 [-0.008-0.135] | 0.084 [0.012-0.227] | 0.014 | 0.134 | 0.017 |
| Lymphoid | IMMATURE B | 0.01 [-0.015-0.117] | 0.052 [0.0-0.145] | 0.084 [0.016-0.179] | 0.457 | 0.033 | 0.097 |
| Lymphoid | LARGE PRE B | 0.015 [-0.013-0.096] | 0.047 [-0.001-0.121] | 0.067 [0.018-0.154] | 0.194 | 0.023 | 0.119 |
| Lymphoid | LATE PRO B | 0.02 [-0.015-0.101] | 0.056 [0.004-0.138] | 0.081 [0.034-0.169] | 0.055 | 0.027 | 0.076 |
| Lymphoid | MATURE B | 0.013 [-0.01-0.1] | 0.06 [0.002-0.125] | 0.084 [0.034-0.163] | 0.231 | 0.013 | 0.038 |
| Lymphoid | NK | 0.024 [-0.006-0.142] | 0.022 [-0.004-0.125] | 0.081 [0.016-0.209] | 0.021 | 0.293 | 0.050 |
| Lymphoid | PLASMA B | 0.005 [-0.009-0.099] | 0.011 [-0.008-0.102] | 0.034 [-0.011-0.158] | 0.874 | 0.749 | 0.718 |
| Lymphoid | PRE PRO B | 0.023 [-0.009-0.14] | 0.041 [-0.006-0.129] | 0.094 [0.026-0.198] | 0.020 | 0.061 | 0.013 |
| Lymphoid | PRO B | 0.011 [-0.02-0.107] | 0.069 [0.008-0.148] | 0.085 [0.041-0.179] | 0.406 | 0.003 | 0.035 |
| Lymphoid | SMALL PRE B | 0.015 [-0.011-0.112] | 0.03 [-0.003-0.124] | 0.066 [0.01-0.169] | 0.150 | 0.153 | 0.179 |
| Lymphoid | TREG | 0.017 [-0.013-0.133] | 0.058 [-0.007-0.141] | 0.09 [0.03-0.189] | 0.122 | 0.025 | 0.045 |

| | | | | | | | |
|---|---|---|---|---|---|---|---|
| Lymphoid | TYPE 1 INNATE T | 0.016 [-0.012-0.142] | 0.065 [-0.009-0.151] | 0.09 [0.017-0.216] | 0.132 | 0.023 | 0.076 |
| Lymphoid | TYPE 3 INNATE T | 0.015 [-0.014-0.115] | 0.045 [-0.011-0.137] | 0.075 [0.01-0.177] | 0.126 | 0.058 | 0.117 |
| MegaK Ery | EARLY ERY | 0.02 [0.001-0.112] | 0.019 [-0.0-0.095] | 0.063 [0.013-0.163] | 0.083 | 0.412 | 0.226 |
| MegaK Ery | EARLY MK MegaK Ery | 0.015 [-0.011-0.116] | 0.049 [-0.005-0.13] | 0.085 [0.026-0.174] | 0.110 | 0.029 | 0.029 |
| MegaK Ery | LATE ERY | 0.013 [-0.014-0.14] | 0.045 [-0.007-0.163] | 0.095 [0.0-0.225] | 0.227 | 0.078 | 0.084 |
| MegaK Ery | LATE MK | 0.007 [-0.006-0.079] | 0.02 [-0.006-0.078] | 0.041 [-0.0-0.118] | 0.672 | 0.353 | 0.503 |
| MegaK Ery | MACROPHAGE ERY | 0.009 [-0.005-0.099] | 0.025 [-0.004-0.108] | 0.058 [0.005-0.157] | 0.529 | 0.306 | 0.382 |
| MegaK Ery | MEMP MegaK Ery | 0.017 [-0.005-0.122] | 0.035 [-0.006-0.119] | 0.076 [0.014-0.174] | 0.075 | 0.109 | 0.060 |
| MegaK Ery | MEP MegaK Ery | 0.018 [-0.005-0.123] | 0.041 [-0.007-0.125] | 0.082 [0.02-0.177] | 0.078 | 0.071 | 0.061 |
| MegaK Ery | MID ERY | 0.006 [-0.008-0.076] | 0.024 [-0.006-0.086] | 0.043 [0.0-0.13] | 0.735 | 0.299 | 0.497 |
| MegaK Ery | VSMC PERI-CYTE III | 0.006 [-0.003-0.062] | 0.011 [-0.002-0.057] | 0.033 [0.003-0.094] | 0.766 | 0.678 | 0.640 |
| MegaK Ery | YS ERY | 0.03 [-0.029-0.2] | 0.041 [-0.018-0.202] | 0.09 [-0.012-0.314] | 0.029 | 0.125 | 0.139 |
| Myeloid | AS DC | 0.007 [-0.012-0.07] | 0.031 [-0.005-0.091] | 0.042 [0.0-0.121] | 0.723 | 0.160 | 0.543 |
| Myeloid | CMP | 0.014 [-0.007-0.101] | 0.037 [-0.006-0.11] | 0.068 [0.011-0.146] | 0.212 | 0.083 | 0.124 |
| Myeloid | DC1 | 0.012 [-0.009-0.098] | 0.045 [-0.006-0.114] | 0.064 [0.012-0.15] | 0.283 | 0.052 | 0.148 |
| Myeloid | DC2 | 0.016 [-0.009-0.123] | 0.031 [-0.004-0.126] | 0.078 [0.011-0.181] | 0.152 | 0.139 | 0.100 |
| Myeloid | DC PROGENI-TOR | 0.017 [-0.014-0.116] | 0.05 [-0.006-0.134] | 0.074 [0.019-0.18] | 0.104 | 0.039 | 0.114 |
| Myeloid | EOSINOPHIL BA-SOPHIL | 0.014 [-0.002-0.099] | 0.018 [-0.001-0.088] | 0.049 [0.004-0.156] | 0.191 | 0.370 | 0.344 |
| Myeloid | GMP | 0.009 [-0.003-0.081] | 0.014 [-0.004-0.079] | 0.038 [0.002-0.122] | 0.537 | 0.602 | 0.573 |
| Myeloid | LANGERHANS CELLS | 0.01 [-0.01-0.12] | 0.012 [-0.009-0.112] | 0.033 [-0.014-0.188] | 0.485 | 0.759 | 0.749 |
| Myeloid | MACROPHAGE IRON RECY-CLING | 0.02 [0.001-0.118] | 0.021 [-0.0-0.095] | 0.059 [0.009-0.177] | 0.042 | 0.295 | 0.177 |
| Myeloid | MACROPHAGE KUPFFER LIKE | 0.011 [-0.004-0.092] | 0.033 [-0.001-0.091] | 0.059 [0.01-0.138] | 0.370 | 0.203 | 0.239 |
| Myeloid | MACROPHAGE LYVE1 HIGH | 0.015 [-0.004-0.112] | 0.048 [-0.003-0.121] | 0.084 [0.022-0.167] | 0.158 | 0.041 | 0.032 |
| Myeloid | MACROPHAGE MHCII HIGH | 0.007 [-0.004-0.088] | 0.01 [-0.005-0.09] | 0.04 [-0.005-0.141] | 0.668 | 0.736 | 0.589 |

| | | | | | | | |
|---|---|---|---|---|---|---|---|
| Myeloid | MACROPHAGE TREM2 | 0.021 [-0.001-0.122] | 0.037 [-0.001-0.114] | 0.079 [0.021-0.176] | 0.057 | 0.132 | 0.093 |
| Myeloid | MAST CELL | 0.019 [-0.008-0.142] | 0.059 [-0.006-0.14] | 0.096 [0.032-0.201] | 0.052 | 0.006 | 0.003 |
| Myeloid | MIGRATORY DC | 0.013 [-0.008-0.098] | 0.031 [-0.002-0.104] | 0.066 [0.011-0.155] | 0.254 | 0.167 | 0.169 |
| Myeloid | MONOCYTE III IL1B | 0.017 [-0.006-0.128] | 0.032 [-0.005-0.127] | 0.082 [0.009-0.188] | 0.102 | 0.162 | 0.079 |
| Myeloid | MONOCYTE II CCR2 | 0.008 [-0.008-0.093] | 0.015 [-0.004-0.097] | 0.046 [-0.006-0.148] | 0.585 | 0.559 | 0.515 |
| Myeloid | MONOCYTE I CXCR4 | 0.011 [-0.004-0.097] | 0.024 [-0.005-0.099] | 0.055 [0.002-0.148] | 0.306 | 0.295 | 0.317 |
| Myeloid | MOP | 0.009 [-0.009-0.09] | 0.028 [-0.009-0.1] | 0.052 [0.003-0.137] | 0.561 | 0.149 | 0.294 |
| Myeloid | MYELOCYTE | 0.011 [-0.015-0.117] | 0.023 [-0.013-0.147] | 0.059 [-0.015-0.199] | 1.000 | 1.000 | 1.000 |
| Myeloid | NEUTROPHIL | 0.015 [-0.021-0.152] | 0.056 [-0.013-0.178] | 0.098 [0.001-0.233] | 0.191 | 0.064 | 0.104 |
| Myeloid | OSTEOCLAST | 0.004 [-0.004-0.065] | 0.005 [-0.004-0.054] | 0.018 [-0.006-0.106] | 0.920 | 0.950 | 0.909 |
| Myeloid | PDC | 0.014 [-0.006-0.111] | 0.025 [0.001-0.111] | 0.075 [0.018-0.166] | 0.135 | 0.217 | 0.048 |
| Myeloid | PRE DC2 | 0.01 [-0.012-0.102] | 0.043 [-0.002-0.122] | 0.069 [0.013-0.16] | 0.442 | 0.056 | 0.126 |
| Myeloid | PROMONOCYTE | 0.014 [-0.008-0.112] | 0.056 [-0.004-0.117] | 0.074 [0.022-0.156] | 0.195 | 0.012 | 0.069 |
| Myeloid | PROMYELOCYTE | 0.007 [-0.005-0.084] | 0.01 [-0.007-0.074] | 0.038 [-0.005-0.119] | 0.719 | 0.770 | 0.626 |

**Supplementary Table 5.** Inferred $\omega_a$ value and 95% confidence interval for each cell lineage posterior distribution in the Adult Atlas.

| Lineage | Cell | $\omega_{aW}$ | $\omega_{aS}$ | $\omega_a$ | Pvalue $\omega_{aW}$ | Pvalue $\omega_{aS}$ | Pvalue $\omega_a$ |
|---|---|---|---|---|---|---|---|
| Bcells | ABCs | 0.011 [-0.004-0.09] | 0.011 [-0.009-0.088] | 0.054 [-0.001-0.139] | 0.481 | 0.623 | 0.376 |
| Bcells | GC B I | 0.011 [-0.012-0.081] | 0.036 [-0.009-0.099] | 0.055 [0.004-0.131] | 0.499 | 0.067 | 0.308 |
| Bcells | GC B II | 0.012 [-0.007-0.115] | 0.023 [-0.013-0.119] | 0.065 [0.005-0.166] | 0.445 | 0.246 | 0.257 |
| Bcells | Memory B cells | 0.016 [-0.008-0.104] | 0.036 [-0.011-0.109] | 0.066 [0.004-0.155] | 0.176 | 0.088 | 0.189 |
| Bcells | Naive B cells | 0.01 [-0.01-0.113] | 0.014 [-0.011-0.096] | 0.039 [-0.016-0.165] | 0.448 | 0.478 | 0.567 |
| Bcells | Plasma cells | 0.013 [-0.005-0.088] | 0.012 [-0.008-0.077] | 0.053 [-0.0-0.121] | 0.305 | 0.482 | 0.267 |
| Bcells | Plasmablasts | 0.008 [-0.011-0.079] | 0.018 [-0.009-0.089] | 0.044 [-0.002-0.118] | 0.633 | 0.265 | 0.394 |
| Bcells | Pre-B | 0.016 [-0.017-0.121] | 0.036 [-0.007-0.152] | 0.083 [0.004-0.194] | 0.219 | 0.144 | 0.168 |
| Bcells | Pro-B | 0.008 [-0.018-0.163] | 0.025 [-0.019-0.187] | 0.075 [-0.021-0.251] | 1.000 | 1.000 | 1.000 |
| Myeloid | Alveolar macrophages | 0.004 [-0.006-0.074] | 0.013 [-0.005-0.075] | 0.038 [-0.001-0.115] | 0.876 | 0.600 | 0.572 |
| Myeloid | Classical monocytes | 0.027 [-0.007-0.136] | 0.035 [-0.006-0.126] | 0.089 [0.013-0.198] | 0.012 | 0.143 | 0.030 |
| Myeloid | DC1 | 0.018 [-0.016-0.113] | 0.069 [0.008-0.145] | 0.1 [0.048-0.183] | 0.081 | 0.004 | 0.010 |
| Myeloid | DC2 | 0.008 [-0.011-0.095] | 0.05 [0.002-0.118] | 0.075 [0.025-0.15] | 0.634 | 0.038 | 0.109 |
| Myeloid | Erythrophagocytic macrophages | 0.009 [-0.009-0.093] | 0.01 [-0.012-0.104] | 0.04 [-0.015-0.149] | 0.602 | 0.857 | 0.689 |
| Myeloid | Intermediate macrophages | 0.01 [-0.009-0.092] | 0.02 [-0.004-0.101] | 0.051 [-0.004-0.146] | 0.450 | 0.364 | 0.427 |
| Myeloid | Intestinal macrophages | 0.011 [-0.004-0.093] | 0.038 [0.005-0.099] | 0.065 [0.016-0.149] | 0.355 | 0.085 | 0.149 |
| Myeloid | Nonclassical monocytes | 0.024 [-0.015-0.137] | 0.036 [-0.008-0.119] | 0.083 [0.01-0.197] | 0.029 | 0.109 | 0.045 |
| Myeloid | migDC | 0.01 [-0.012-0.107] | 0.058 [0.001-0.134] | 0.074 [0.017-0.171] | 0.423 | 0.038 | 0.173 |
| Tcells | ILC3 | 0.017 [-0.003-0.117] | 0.021 [-0.003-0.098] | 0.068 [0.007-0.17] | 0.126 | 0.220 | 0.070 |
| Tcells | MAIT | 0.036 [-0.024-0.207] | 0.045 [-0.02-0.182] | 0.102 [-0.002-0.296] | 0.009 | 0.047 | 0.046 |
| Tcells | NK CD16+ | 0.013 [-0.007-0.114] | 0.007 [-0.011-0.087] | 0.03 [-0.012-0.179] | 0.333 | 0.662 | 0.639 |
| Tcells | NK CD56bright CD16- | 0.04 [-0.001-0.177] | 0.031 [-0.015-0.117] | 0.089 [0.017-0.256] | 0.002 | 0.053 | 0.006 |
| Tcells | T CD4 CD8 | 0.016 [-0.036-0.186] | 0.046 [-0.031-0.232] | 0.1 [-0.031-0.305] | 1.000 | 1.000 | 1.000 |
| Tcells | Teffector EM CD4 | 0.012 [-0.01-0.106] | 0.015 [-0.01-0.104] | 0.052 [-0.012-0.159] | 0.363 | 0.462 | 0.395 |
| Tcells | Tem emra CD8 | 0.008 [-0.005-0.089] | 0.01 [-0.007-0.084] | 0.039 [-0.005-0.131] | 0.597 | 0.574 | 0.496 |
| Tcells | Tfh | 0.007 [-0.006-0.089] | 0.014 [-0.007-0.097] | 0.043 [-0.006-0.139] | 0.666 | 0.492 | 0.517 |
| Tcells | Tgd CRTAM+ | 0.009 [-0.009-0.1] | 0.012 [-0.009-0.079] | 0.03 [-0.012-0.15] | 0.553 | 0.573 | 0.682 |
| Tcells | Tnaive CM CD4 | 0.009 [-0.005-0.084] | 0.006 [-0.008-0.066] | 0.018 [-0.009-0.122] | 0.579 | 0.841 | 0.903 |

| | | | | | | | |
|---|---|---|---|---|---|---|---|
| Tcells | Tnaive CM CD4 activated | 0.013 [0.0-0.087] | 0.012 [-0.009-0.082] | 0.044 [-0.001-0.128] | 0.352 | 0.525 | 0.463 |
| Tcells | Tnaive CM CD8 | 0.005 [-0.005-0.059] | 0.004 [-0.005-0.047] | 0.013 [-0.008-0.092] | 0.828 | 0.954 | 0.968 |
| Tcells | Tregs | 0.014 [-0.01-0.129] | 0.016 [-0.008-0.118] | 0.057 [-0.008-0.182] | 0.234 | 0.355 | 0.268 |
| Tcells | Trm Tgd | 0.018 [-0.001-0.12] | 0.015 [-0.008-0.094] | 0.062 [0.007-0.172] | 0.123 | 0.289 | 0.088 |
| Tcells | Trm Th1 Th17 | 0.053 [-0.056-0.304] | 0.05 [-0.036-0.247] | 0.148 [0.031-0.407] | 0.002 | 0.042 | 0.001 |
| Tcells | Trm em CD8 | 0.011 [-0.02-0.149] | 0.054 [-0.022-0.175] | 0.081 [-0.002-0.222] | 0.307 | 0.041 | 0.154 |
| Tcells | Trm gut CD8 | 0.015 [-0.008-0.134] | 0.032 [-0.013-0.121] | 0.073 [0.0-0.187] | 0.231 | 0.096 | 0.087 |

**Supplementary Table 6.** Inferred $\omega_a$ value and 95% confidence interval on iPSC lines differentiated into Macrophages and exposed to 10 stimuli that emulate responses to pro/anti-inflammatory cytokines and bacterial or viral infections at two different timepoints.

| Stimuli | Time | $\omega_{aW}$ | $\omega_{aS}$ | $\omega_a$ | Pvalue $\omega_{aW}$ | Pvalue $\omega_{aS}$ | Pvalue $\omega_a$ |
|---|---|---|---|---|---|---|---|
| CIL | Early | 0.019 [-0.01-0.132] | 0.071 [-0.004-0.144] | 0.102 [0.041-0.193] | 0.021 | 0.005 | 0.003 |
| CIL | Inter | 0.007 [-0.005-0.081] | 0.011 [-0.004-0.07] | 0.032 [-0.001-0.12] | 0.623 | 0.689 | 0.671 |
| CIL | Late | 0.007 [-0.006-0.084] | 0.02 [-0.005-0.088] | 0.039 [-0.001-0.127] | 0.474 | 0.349 | 0.346 |
| IFNB | Early | 0.007 [-0.009-0.097] | 0.01 [-0.006-0.079] | 0.025 [-0.012-0.152] | 0.822 | 0.824 | 0.844 |
| IFNB | Inter | 0.01 [-0.009-0.09] | 0.031 [-0.005-0.099] | 0.055 [0.005-0.134] | 0.334 | 0.126 | 0.205 |
| IFNB | Late | 0.006 [-0.007-0.073] | 0.025 [-0.0-0.085] | 0.046 [0.006-0.116] | 0.522 | 0.173 | 0.267 |
| IFNG | Early | 0.025 [-0.014-0.172] | 0.055 [-0.011-0.169] | 0.118 [0.022-0.251] | 0.002 | 0.041 | 0.003 |
| IFNG | Inter | 0.007 [-0.007-0.085] | 0.004 [-0.007-0.055] | 0.014 [-0.011-0.122] | 0.968 | 0.984 | 0.984 |
| IFNG | Late | 0.004 [-0.008-0.083] | 0.013 [-0.004-0.091] | 0.045 [-0.002-0.129] | 0.593 | 0.415 | 0.466 |
| IL4 | Early | 0.008 [-0.013-0.11] | 0.015 [-0.01-0.13] | 0.057 [-0.013-0.187] | 0.615 | 0.441 | 0.496 |
| IL4 | Inter | 0.02 [-0.014-0.128] | 0.042 [-0.009-0.142] | 0.08 [0.012-0.185] | 0.031 | 0.013 | 0.021 |
| IL4 | Late | 0.013 [-0.01-0.122] | 0.032 [-0.005-0.134] | 0.064 [-0.006-0.197] | 0.764 | 0.575 | 0.531 |
| LIL10 | Early | 0.01 [-0.008-0.089] | 0.024 [-0.006-0.101] | 0.057 [0.008-0.138] | 0.400 | 0.146 | 0.162 |
| LIL10 | Inter | 0.013 [-0.005-0.109] | 0.02 [-0.003-0.097] | 0.059 [0.001-0.157] | 0.207 | 0.286 | 0.230 |
| LIL10 | Late | 0.006 [-0.006-0.082] | 0.021 [-0.0-0.097] | 0.048 [0.003-0.137] | 0.494 | 0.158 | 0.306 |
| MBP | Early | 0.009 [-0.015-0.123] | 0.034 [-0.002-0.141] | 0.073 [0.002-0.182] | 0.182 | 0.047 | 0.086 |
| MBP | Inter | 0.008 [-0.011-0.107] | 0.018 [-0.007-0.119] | 0.051 [-0.011-0.168] | 0.325 | 0.286 | 0.218 |
| MBP | Late | 0.009 [-0.009-0.102] | 0.013 [-0.008-0.097] | 0.039 [-0.012-0.163] | 0.937 | 0.907 | 0.942 |
| P3C | Early | 0.012 [-0.006-0.098] | 0.043 [-0.001-0.112] | 0.07 [0.019-0.157] | 0.225 | 0.056 | 0.079 |
| P3C | Inter | 0.006 [-0.009-0.1] | 0.011 [-0.009-0.081] | 0.023 [-0.014-0.164] | 0.695 | 0.787 | 0.798 |
| P3C | Late | 0.013 [-0.014-0.102] | 0.041 [-0.004-0.133] | 0.073 [0.011-0.161] | 0.818 | 0.135 | 0.189 |
| PIC | Early | 0.017 [-0.008-0.136] | 0.02 [-0.01-0.153] | 0.048 [-0.008-0.238] | 0.668 | 0.798 | 0.707 |
| PIC | Inter | 0.009 [-0.003-0.091] | 0.023 [-0.003-0.095] | 0.057 [0.005-0.138] | 0.252 | 0.224 | 0.145 |
| PIC | Late | 0.009 [-0.006-0.079] | 0.02 [-0.005-0.079] | 0.043 [0.001-0.117] | 0.272 | 0.267 | 0.328 |
| R484 | Early | 0.013 [-0.01-0.111] | 0.041 [-0.003-0.124] | 0.08 [0.017-0.172] | 0.110 | 0.038 | 0.040 |
| R484 | Inter | 0.005 [-0.009-0.096] | 0.008 [-0.008-0.082] | 0.031 [-0.013-0.143] | 0.726 | 0.779 | 0.681 |
| R484 | Late | 0.008 [-0.009-0.097] | 0.025 [-0.005-0.109] | 0.063 [0.001-0.144] | 0.402 | 0.237 | 0.187 |
| all | Early | 0.009 [-0.006-0.092] | 0.031 [0.001-0.095] | 0.06 [0.01-0.145] | 0.170 | 0.147 | 0.131 |
| all | Inter | 0.014 [-0.006-0.131] | 0.028 [-0.003-0.127] | 0.076 [0.004-0.196] | 0.050 | 0.095 | 0.099 |
| all | Late | 0.007 [-0.005-0.073] | 0.01 [-0.005-0.065] | 0.029 [-0.005-0.112] | 0.656 | 0.634 | 0.689 |
| sLPS | Early | 0.007 [-0.005-0.077] | 0.021 [-0.002-0.08] | 0.049 [0.007-0.118] | 0.243 | 0.238 | 0.225 |
| sLPS | Inter | 0.009 [-0.007-0.095] | 0.022 [-0.003-0.094] | 0.049 [0.003-0.141] | 0.451 | 0.396 | 0.376 |
| sLPS | Late | 0.016 [-0.006-0.108] | 0.037 [0.0-0.121] | 0.074 [0.014-0.174] | 0.060 | 0.081 | 0.059 |

**Supplementary Table 7.** ABC-MK prior distributions

| $\alpha$ | Weak adaptive selection coefficient | Strong adaptive selection coefficient | Mean deleterious selection coefficient | Gamma distribution shape for deleterious alleles | Deleterious selection coefficient driving BGS | BGS values (McVicker et al.[91] B) |
|---|---|---|---|---|---|---|
| $[0.0, 0.9]$ | $[1, 10]$ | $[200, 2000]$ | $[-1000, -200]$ | $0.184 \cdot 2^{[-2,2]}$ | $[-1000, -500]$ | $[0.1, 0.999]$ |

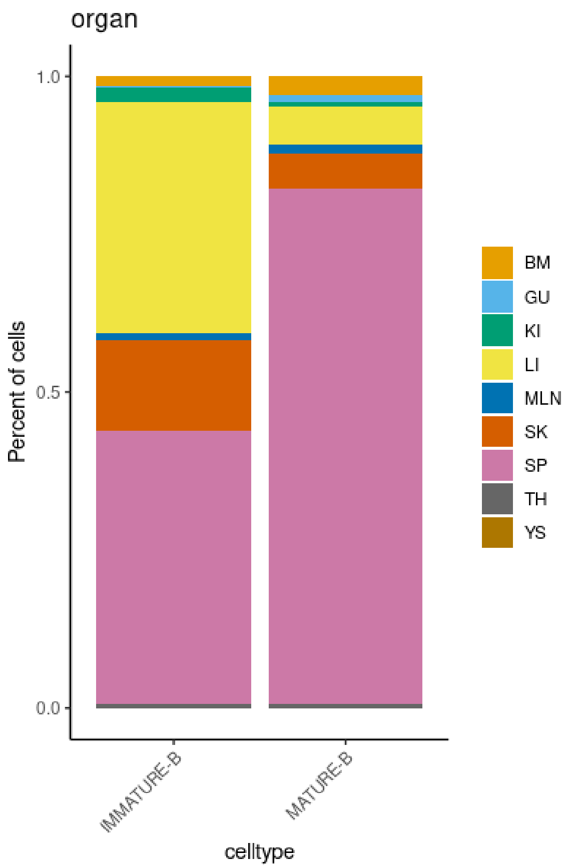


**Supplementary Figure 1.** Distribution across organs of Immature and Mature B cells in the Developmental Cell Atlas. BM, bone marrow; GU, gut; KI, kidney; LI, liver; MLN, mesenteric lymph node; SK, skin; SP, spleen; TH, thymus; YS, yolk sac;

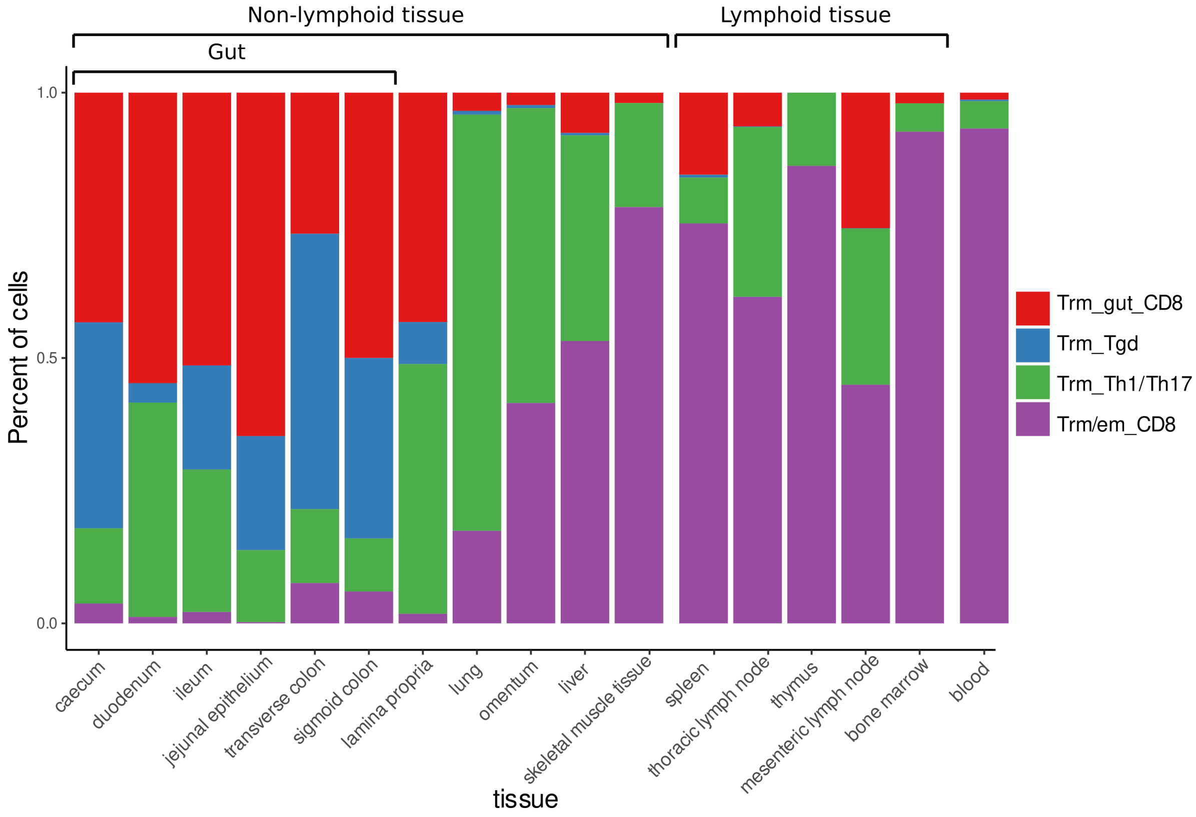


**Supplementary Figure 2.** Distribution of T resident memory (Trm) cells across tissues

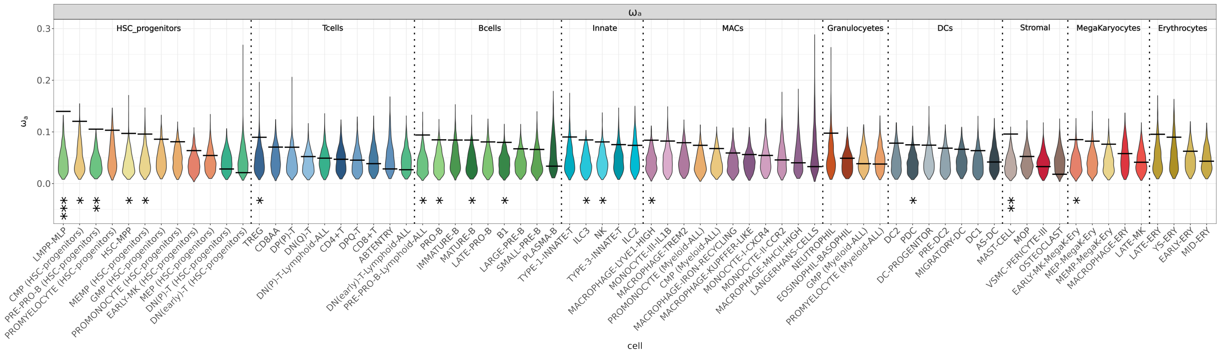


**Supplementary Figure 3.** Protein adaptation in the Developmental Atlas. $\omega_a$ value of each cell type as a horizontal bar and the $\omega_a$ value distribution of its control bootstrap dataset ($n = 1000$) as a violin plot. The asterisks highlight the cell types with a significant high $\omega_a$ (* p-value $< 0.05$, ** p-value $< 0.01$).

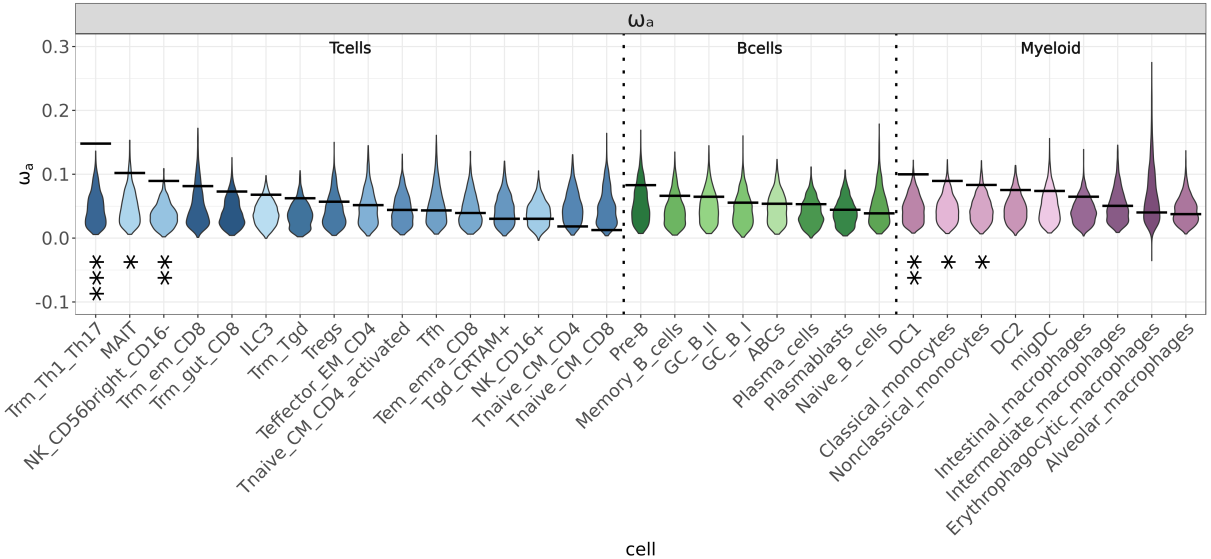


**Supplementary Figure 4.** Protein adaptation in the Adult Atlas. $\omega_a$ value of each cell type as a horizontal bar and the $\omega_a$ value distribution of its control bootstrap dataset ($n = 1000$) as a violin plot. The asterisks highlight the cell types with a significant high $\omega_a$ (* p-value $< 0.05$, ** p-value $< 0.01$).

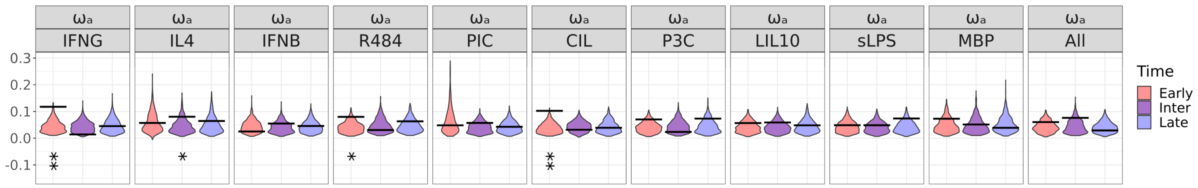


**Supplementary Figure 5.** Protein adaptation in Activated Macrophages. $\omega_a$ estimation for each stimulus and time-specific responses categorised as "early" (exclusively after 6h), "late" (exclusively after 24h) or "sustained" (found at both time points). For each stimulus and time-specific response we show the inferred $\omega_a$ value (horizontal bar) and the distribution of the control bootstrap dataset (n=1000) as a violin plot. The asterisks highlight the responses with a significant high $\alpha$ (* p-value $<$ 0.05, ** p-value $<$ 0.01).
